# *Agrobacterium tumefaciens* divisome proteins regulate the transition from polar growth to cell division

**DOI:** 10.1101/412759

**Authors:** Matthew Howell, Alena Aliashkevich, Kousik Sundararajan, Jeremy J. Daniel, Patrick J. Lariviere, Erin D. Goley, Felipe Cava, Pamela J.B. Brown

## Abstract

The mechanisms that restrict peptidoglycan biosynthesis to the pole during elongation and re-direct peptidoglycan biosynthesis to mid-cell during cell division in polar-growing Alphaproteobacteria are largely unknown. Here, we demonstrate that although two of the three FtsZ homologs localize to mid-cell, exhibit GTPase activity and form co-polymers, only one, FtsZ_AT_, is required for cell division. We find that FtsZ_AT_ is required not only for constriction and cell separation, but also for the termination of polar growth and regulation of peptidoglycan synthesis at mid-cell. Depletion of FtsZ in *A. tumefaciens* causes a striking phenotype: cells are extensively branched and accumulate growth active poles through tip splitting events. When cell division is blocked at a later stage, polar growth is terminated and ectopic growth poles emerge from mid-cell. Overall, this work suggests that *A. tumefaciens* FtsZ makes distinct contributions to the regulation of polar growth and cell division.

## Introduction

The spatial and temporal regulation of cell division is a vital process across bacterial species with implications in the development of antimicrobial therapies [1]. The cell division process must coordinate membrane invagination(s), peptidoglycan (PG) biosynthesis and remodeling, and the physical separation of the two daughter cells, all while maintaining cellular integrity. Furthermore, cell division must be precisely regulated to be orchestrated with other key cell cycle processes including cell elongation, DNA replication, and chromosome segregation to ensure that each daughter cell is of sufficient size and contains a complete genome [2, 3].

To initiate bacterial cell division, the tubulin-like GTPase, FtsZ, polymerizes and forms a discontinuous ring-like structure at the future site of cell division [4-10]. The presence of FtsZ at mid-cell leads to the recruitment of many proteins that function in cell division, collectively called the divisome [11-14]. The divisome includes cell wall biosynthesis proteins, such as the penicillin-binding protein, PBP3, and FtsW, which contribute to PG biosynthesis and remodeling necessary to form new poles in daughter cells [11]. Once the divisome is fully assembled, FtsZ filaments treadmill along the circumference of the mid-cell, driving the Z-ring constriction [9, 10]. The movement of FtsZ filaments is correlated with the movement of enzymes that function in septal PG biogenesis. These finding are consistent with the notion that FtsZ not only recruits enzymes that function in PG biogenesis to mid-cell but also regulates their activities to promote proper cell wall biogenesis [15-17].

In most rod-shaped model organisms used to study cell division, a block in cell division leads to the production of long, smooth filamentous cells. This phenotype suggests that assembly or activation of some divisome components is necessary not only to enable the cells to divide but also to stop cellular elongation. Indeed, in *Escherichia coli*, FtsZ (along with the Z-ring stabilizing proteins FtsA, ZipA, and ZapA) has been proposed to have an early function in the switch from lateral PG biogenesis to mid-cell PG biosynthesis [18]. Following maturation of the divisome by recruitment of additional PG remodeling enzymes and cell division proteins, PG biosynthesis is coordinated with membrane invagination, enabling cells to constrict and separate [19].

Conversely to e.g. *E. coli*, polar growing rods in the alphaproteobacterial clade Rhizobiales exhibit branched morphologies when cell division is blocked [20-27]. Examination of the cell morphologies resulting from the block in cell division suggests that different types of branched morphologies arise [28]. Drug treatments that block DNA replication cause an early block in cell division, resulting in a “Y” morphology in which the branches are formed from existing growth poles [25, 26]. In contrast, antibiotics that target PBP3 cause mid-cell bulges and branches with some cells adopting a “T” or “+” morphologies [25, 27]. These observations suggest that polar-like PG synthesis is redirected to mid-cell when cell division is blocked at a later stage. The manifestation of two distinct phenotypes during early and late blocks in cell division suggests that divisome assembly and activation may contribute to termination of polar growth, onset of mid-cell PG biosynthesis, cell constriction, and ultimately cell separation.

In *Agrobacterium tumefaciens,* homologs of FtsZ and FtsA fused to fluorescent proteins localize at the growth pole during elongation and at mid-cell during division [27, 29, 30]. FtsZ was found to arrive at mid-cell considerably earlier than FtsA [30], indicating that FtsZ may be able to initiate Z-ring formation prior to FtsA recruitment to the divisome. This observation is consistent with the described order of divisome assembly in *Caulobacter crescentus* [31] and suggests that a distinctive time-dependent role of these proteins in cell division.

Here, we take advantage of the ability to deplete essential proteins in *A. tumefaciens* [32] to explore the function of cell division proteins FtsZ, FtsA, and FtsW in a polar growing alphaproteobacterium. Although the genome of *A. tumefaciens* encodes three Fts*Z* homologs, we find that only one, henceforth referred to as FtsZ_AT,_ is essential for cell survival. FtsZ_AT_ is required to recruit division proteins to mid-cell and likely regulates the activity of PG biosynthesis enzymes at mid-cell. In the absence of FtsZ_AT_, cells not only fail to divide but are also unable to terminate polar growth. Depletion of either FtsA or FtsW also causes a block in cell division, but unlike FtsZ_AT_ depletion, growth at the poles is halted and instead, polar-like PG synthesis is redirected to mid-cell. These observations suggest that only FtsZ is required to terminate polar growth and initiate cell division-specific PG biosynthesis at mid-cell, whereas FtsZ, FtsA, and FtsW are exclusively required for cell division. Together these findings suggest that *A. tumefaciens* uses sequential regulation of cell division, a theme that is broadly conserved in bacteria.

## Results and Discussion

### FtsZ_AT_ is required for cell division and termination of polar growth

*Agrobacterium tumefaciens* contains three homologs of *Escherichia coli’s* FtsZ, Atu_2086, Atu_4673, and Atu_4215 (Figure 1A) [27]. *E. coli* FtsZ is comprised of three regions: the conserved N-terminal tubulin-like GTPase domain, a C-terminal linker (CTL), and a conserved C-terminal peptide (CTP), which anchors FtsZ to the membrane via interactions with FtsA [33]. Atu_2086 contains each of these domains out of which the GTPase domain and CTP share 52% and 67% identity to their respective domain in *E. coli* FtsZ, whereas the CTL is extended in length [27]. The gene encoding Atu_2086 is found in a putative operon with genes encoding DdlB, FtsQ, FtsA [34, 35] and is predicted to be essential for cell survival based on saturating transposon mutagenesis [36]. Atu_2086 localizes to mid-cell in wildtype (WT) pre-divisional cells (Figure 1B) [27, 29]; consistent with a role in cell division. Atu_4673 (called FtsZ_1_; consistent with the genome annotation) contains a complete GTPase domain with 49% identity to tubulin domain of *E. coli* FtsZ but lacks both the CTL and CTP [27]. Although Atu_4673 is not predicted to be required for cell survival based on saturating transposon mutagenesis [36], it localizes to mid-cell in pre-divisional cells, suggesting a possible role in cell division (Figure 1B). Atu_4215 (termed FtsZ_3_ in this work) contains a partial GTPase domain with 48% identity to the N-terminal portion of the *E. coli* FtsZ tubulin domain and lacks both the CTL and CTP [27]. FtsZ_3_ is not essential for survival of *A. tumefaciens* based on saturating transposon mutagenesis [36] and exhibits a diffuse localization pattern (Figure 1B). Together, these data suggest that Atu_2086 is the canonical FtsZ protein required for cell division, and this protein will be referred to as FtsZ_AT_ throughout this work (although it is annotated as FtsZ_2_ in the *A. tumefaciens* C58 genome [34, 35]).

**Figure 1.**
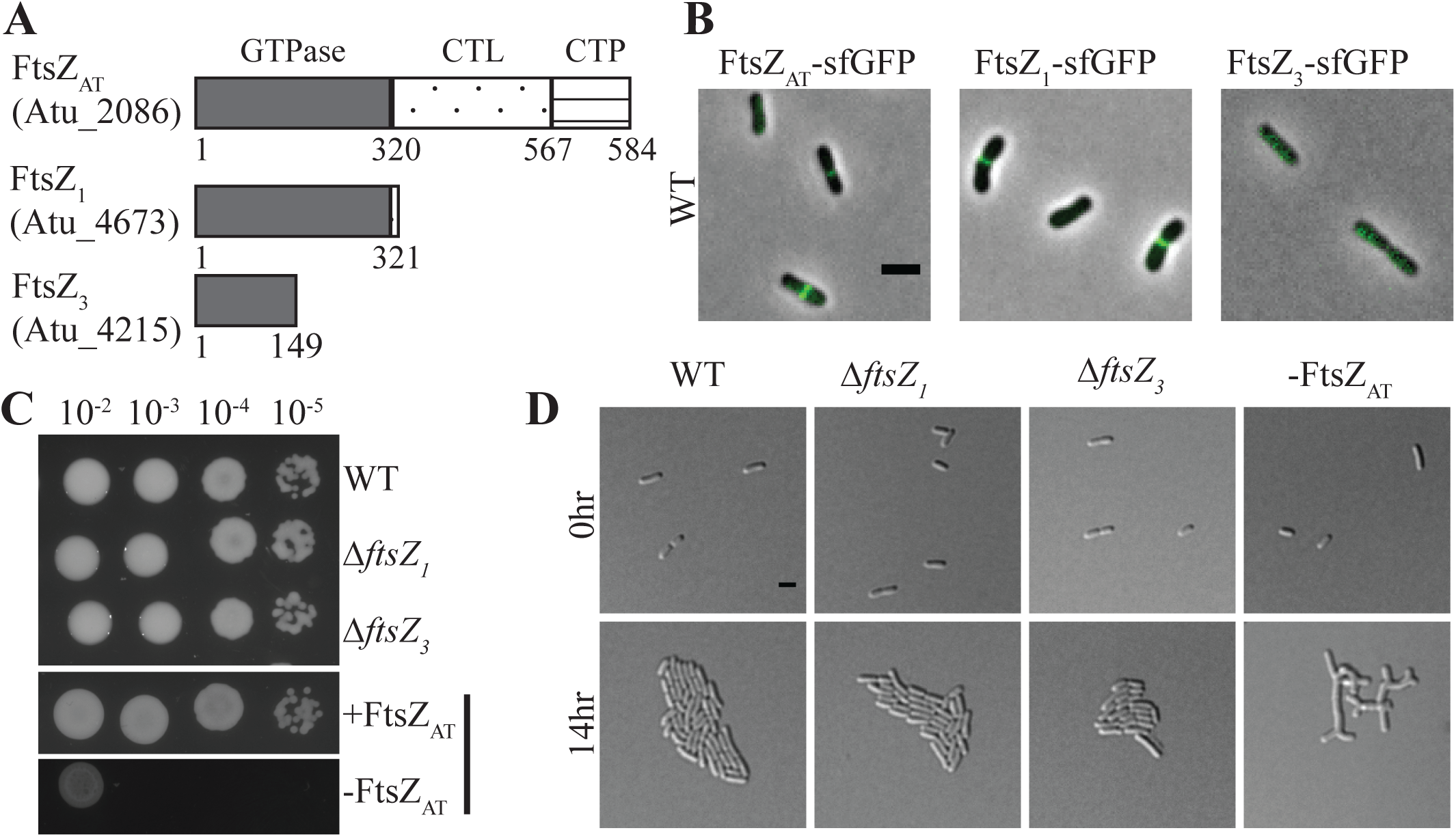
Characterization of FtsZ homologs in *A. tumefaciens*. A) Domain schematic of FtsZ homologs in *A. tumefaciens*. Note that domains are not drawn to scale. B) Representative image of localization patterns for each homolog. FtsZ_AT_-sfGFP and FtsZ_1_-sfGFP show mid-cell ring formation while FtsZ_3_-sfGFP fails to localize. C) Cell viability is shown by spotting serial dilutions. Δ*ftsZ*_*1*_, Δ*ftsZ*_*3*_, and +*ftsZ*_*AT*_ have similar viability to WT, while -*ftsZ*_*AT*_ displays a drastic decrease in viability. D) Cell morphology and microcolony formation of Δ*ftsZ*_*1*_ and Δ*ftsZ*_*3*_ are similar to WT, while -*ftsZ*_*AT*_ results in long branched cells that fail to divide. All scale bars are set to 2 µm.

To characterize the function of each FtsZ homolog, we constructed deletions of *ftsZ*_*1*_ and *ftsZ*_*3*_ and a depletion strain of *ftsZ*_*AT.*_ Since we were unable to construct a deletion of *ftsZ*_*AT*_, we used a depletion strategy in which *ftsZ*_*AT*_ is present as a single copy under the control of an isopropyl β-D-1-thiogalactopyranoside (IPTG) inducible promoter at a neutral site in the chromosome [21, 32]. Using western blot analysis, we have confirmed the depletion of FtsZ_AT_ in the absence of IPTG (Figure 1-Figure Supplement 1A).

Deletion of *ftsZ*_*1*_ or *ftsZ*_*3*_ does not impact cell viability (Figure 1C), cell morphology (Figure 1D; Table 1; Figure 1-Figure Supplement 1B), microcolony formation (Figure 1D), constriction rate or position (Table 1) when compared to WT cells. Similarly, when FtsZ_AT_ is expressed in the depletion strain (labeled in Figures as +FtsZ_AT_) the cells remain viable (Figure 1C), are similar in size to WT cells (Table 1), properly position constrictions (Table 1), and form microcolonies (Figure 1D). In contrast, depletion of FtsZ_AT_ (labeled in Figures as –FtsZ_AT_) causes a marked decrease in cell viability (Figure 1C) and triggers the formation of large cells with complex branched morphologies (Table 1; Figure 1D). To quantify changes in morphology during depletion of FtsZ_AT_, the cell area of at least 100 cells was calculated based on phase contrast images of cells acquired immediately after removal of the inducer (-FtsZ_AT_ 0 h), 8 h after removal of the inducer (-FtsZ_AT_ 8 h), and 14 h after removal of the inducer (-FtsZ_AT_ 14 h) (Table 1, Figure 1-Figure Supplement 1C). Initially, the FtsZ_AT_ depleted cells are similar to WT in cell size, but after 8 h of FtsZ_AT_ depletion the cell area has nearly doubled (Table 1, Figure 1-Figure Supplement 1C). Within 14 h of FtsZ_AT_ depletion, the average cell area has dramatically increased (Table 1, Figure 1-Figure Supplement 1C). Together, these results demonstrate that only the FtsZ_AT_ homolog is required for proper cell growth and division.

**Table 1.** Quantitation of cell size and constriction of *ftsZ* mutants

|  |  | **Average**<br>**Cell Length<sup>a</sup>**<br>**($\mu\text{m}$ +/- SD<sup>b</sup>)** | **Average**<br>**Cell Area<sup>a</sup>**<br>**($\mu\text{m}^2$ +/- SD)** | **Average**<br>**Constriction Rate<sup>c</sup>**<br>**(nm/min +/- SD)** | **Relative**<br>**Constriction**<br>**Position<sup>d</sup>**<br>**+/- SD** |
| --- | --- | --- | --- | --- | --- |
|  | WT | 2.31 +/- .50 | 1.66 +/- .35 | 6.82 +/- 3.19 | .49 +/- .05 |
| | $\Delta ftsZ_1$ | 2.25 +/- .49 | 1.52 +/- .33 | 6.99 +/- 3.58 | .46 +/- .05 |
| | $\Delta ftsZ_3$ | 2.24 +/- .47 | 1.44 +/- .30 | 6.77 +/- 2.77 | .46 +/- .05 |
| | $\Delta ftsZ_1$<br>$\Delta ftsZ_3$ | 2.25 +/- .51 | 1.47 +/- .34 | 6.61 +/- 3.75 | .46 +/- .04 |
| *ftsZ<sub>AT</sub>* depletion | -<br>FtsZ<sub>AT</sub><br>0 h | 2.71 +/- .70 | 1.56 +/- .39 | 6.38 +/- 2.81 | .49 +/- .07 |
|  | -<br>FtsZ<sub>AT</sub><br>8 h | ND<sup>e</sup> | 2.95 +/- 1.12 | ND | ND |
|  | -<br>FtsZ<sub>AT</sub><br>14 h | ND | 11.37 +/- 4.69 | ND | ND |

<sup>a</sup>At least 100 cells were used to quantify the cell length and area for each strain
<sup>b</sup>SD – standard deviation.
<sup>c</sup>At least 30 cells were used to quantify the constriction rates for each strain
<sup>d</sup>Relative constriction position for at least 40 cells is shown. A value of 0 corresponds to the new pole, 0.5 corresponds to mid-cell, and a value of 1 corresponds to the old pole.
<sup>e</sup>ND – not determined.

### Deletion of *ftsZ*_*1*_ and *ftsZ*_*3*_ does not change the FtsZ_AT_ depletion phenotype

Since the *ftsZ*_*1*_ and *ftsZ*_*3*_ single deletions do not have an obvious impact on cell morphology, growth, or division, we constructed double and triple mutants to determine if there is an increasing effect when removing multiple *ftsZ* homologs. Double deletion of *ftsZ*_*1*_ and *ftsZ*_*3*_ does not cause a decrease in cell viability (Figure 2A, top panel), cell morphology (Table 1), or microcolony formation (Figure 2A, bottom panel). Furthermore, *ΔftsZ*_*1*_ *ΔftsZ*_*3*_ cells properly place constrictions and have an average constriction rate similar to WT (Table 1). Next, we introduced the *ΔftsZ*_*1*_, *ΔftsZ*_*3*_, and *ΔftsZ*_*1*_ *ΔftsZ*_*3*_ mutations into the *ftsZ*_*AT*_ depletion strain to determine if loss of multiple *ftsZ* homologs further aggravated the *ftsZ*_*AT*_ depletion phenotypes. The combination of the *ftsZ*_*AT*_ depletion strain with *ΔftsZ*_*1*_, *ΔftsZ*_*3*_, or *ΔftsZ*_*1*_ *ΔftsZ*_*3*_ mutations did not result in a further decrease in cell viability (Figure 2B, top panel) or a worsening of cell morphology (Figure 2B, bottom panel) when compared to FtsZ_AT_ depletion alone. Together, these results suggest that the FtsZ_1_ and FtsZ_3_ homologs do not have a major impact on cell division under the conditions tested.

**Figure 2.**
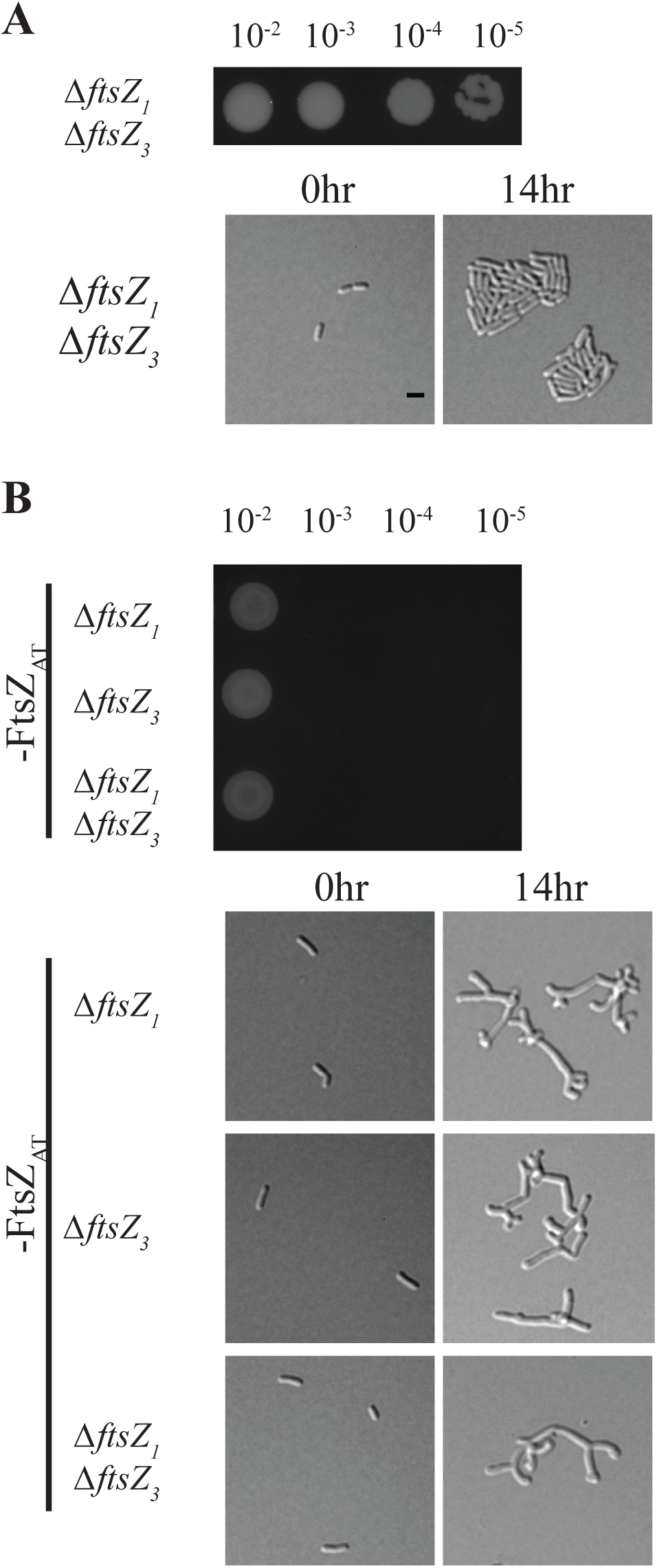
Deletion of multiple *ftsZ* homologs does not yield an additive effect. A) Cell viability (top) and morphology (bottom) of the double mutant Δ*ftsZ*_*1*_Δ*ftsZ*_*3*_ is indistinguishable from WT. B) Cell viability (top) and morphology (bottom) of Δ*ftsZ*_*1*_, Δ*ftsZ*_*3*_, or Δ*ftsZ*_*1*_Δ*ftsZ*_*3*_ during FtsZ_AT_ depletion are indistinguishable from FtsZ_AT_ depletion alone. All scale bars are set to 2 µm. Black bar denotes *ftsZ*_*AT*_ depletion strain background.

*ftsZ* gene duplications have occurred independently in several alphaproteobacterial lineages and in chloroplasts and some mitochondria [37]. In most of the cases that have been studied, one FtsZ homolog plays a canonical role in cell or organelle division while the other plays a regulatory or specialized role. However, little is known about the roles of multiple ftsZs in certain alphaproteobacteria species. In both *Rhizobium meliloti* and *Magnetospirillum gryphiswaldense*, one of the FtsZs (containing a CTL and CTP similar to FtsZ_AT_) is essential and the other (truncated after the GTPase domain similar to FtsZ_1_) is dispensable [38, 39]. In the case of *M. gryphiswaldense*, the truncated *ftsZ* is dispensable for division but important for biomineralization in this magnetotactic species under certain growth conditions. Similarly, it is possible that FtsZ_1_ or FtsZ_3_ may have important contributions to cell growth or division of *A. tumefaciens* in different environments as e.g. in its plant-associated life-style.

### FtsZ_1_ requires FtsZ_AT_ to localize to mid-cell and to polymerize *in vitro*

Since FtsZ_1_ localizes to mid-cell (Figure 1B), we hypothesized that FtsZ_1_ may be a nonessential divisome component. To test this, we examined the localization of FtsZ_1_-sfGFP in both WT and the *ftsZ*_*AT*_ depletion strain (Figure 3). In WT and FtsZ_AT_ induced cells, FtsZ_1_-sfGFP does not localize in newborn cells but forms FtsZ-like rings at the future site of division in pre-divisional cells (Figure 3A, top and middle panel). This Z-like ring constricts to form a single focus in dividing cells. These observations suggest that FtsZ_1_ may be a divisome component despite the absence of a cell division phenotype in the Δ*ftsZ*_*1*_ strain. To explore the possibility of interactions arising due to the loss of FtsZ_1_ and FtsZ_AT_, we next visualized FtsZ_1_-sfGFP localization during the depletion of FtsZ_AT_ (Figure 3A, bottom panel). We pre-depleted FtsZ_AT_ for 4 h in liquid to avoid cell crowding caused by division events prior to sufficient FtsZ_AT_ depletion. Early during the depletion of FtsZ_AT_, FtsZ_1_-sfGFP localizes in a FtsZ-like ring near mid-cell. However, as the FtsZ_AT_ depletion continues, FtsZ_1_-sfGFP rings and foci progressively fade away, demonstrating that localization of FtsZ_1_-sfGFP to mid-cell requires the presence of FtsZ_AT_.

**Figure 3.**
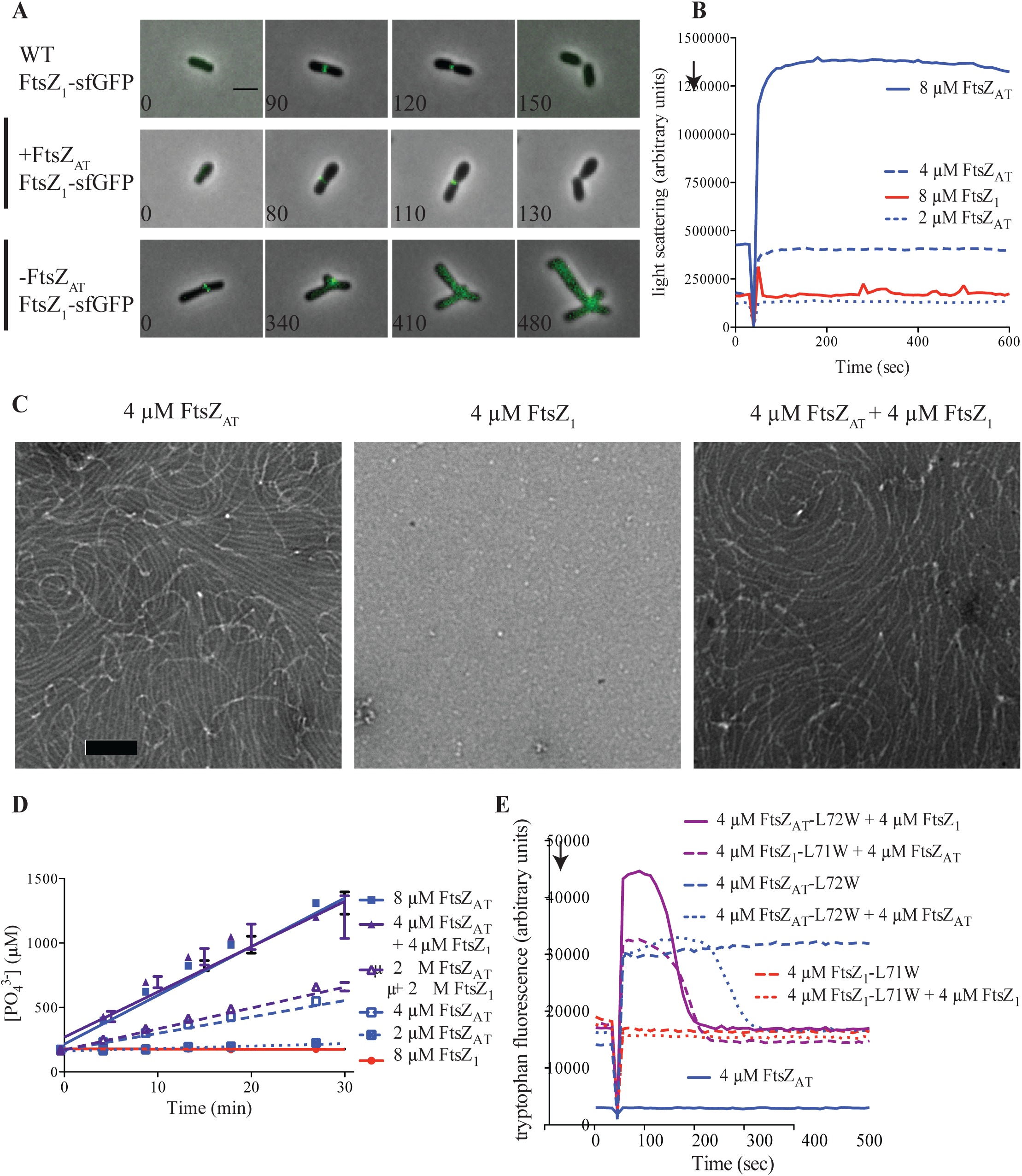
FtsZ_1_ requires FtsZ_AT_ to polymerize *in vitro* and to localize in cells. A) FtsZ_1_-sfGFP forms midcell rings which constrict in WT and when *ftsZ*_*AT*_ is induced. FtsZ_1_-sfGFP fails to constrict early rings and disassembles during FtsZ_AT_ depletion. Scale bar is set to 2 µm. B) Light scattering over time for purified proteins at the indicated concentrations. FtsZ_AT_ polymerizes at concentrations above 2 µM, but FtsZ_1_ does not polymerize. GTP (2 mM) was added where indicated by the arrow to induce polymerization. Experiments were performed in triplicate and mean curves are shown. C) Negative stain TEM of the indicated proteins. Co-polymers of FtsZ_AT_ and FtsZ_1_ are indistinguishable from FtsZ_AT_ polymers. Scale bar is set to 100 nm. D) Inorganic phosphate concentration in solution over time in the presence of the indicated proteins and protein concentrations. Co-polymers of FtsZ_AT_ and FtsZ_1_ consume GTP at the same rate as FtsZ_AT_ homopolymers at equivalent total FtsZ concentration. Reactions were performed in triplicate and mean ± standard error is plotted. E) Tryptophan fluorescence over time for the indicated proteins. FtsZ_1_-L71W (red) shows no polymerization (Trp fluorescence) on its own, but can co-polymerize with added FtsZ_AT_ (purple). GTP (50 µM) was added where indicated by the arrow to induce polymerization. Experiments were performed in triplicate and representative curves are shown.

Since FtsZ_1_ is recruited to mid-cell by FtsZ_AT_, we hypothesized that FtsZ_AT_ and FtsZ_1_ may form co-polymers. To first test the ability of FtsZ_AT_ and FtsZ_1_ to independently form polymers, each protein was purified and subjected to polymerization studies. Right angle light scattering assays of wildtype FtsZ_AT_ revealed that this protein exhibits a GTP-dependent increase in light scattering at concentrations above 2 µm, consistent with its polymerization (Figure 3B, blue lines). Negative stain transmission electron microscopy (TEM) confirmed that FtsZ_AT_ forms gently curved protofilaments in the presence of GTP (Figure 3C, left panel) and it rapidly releases inorganic phosphate suggesting that GTP is hydrolyzed (Figure 3D, blue lines; 4.7 ± 0.2 GTP min^-1^ FtsZ^-1^ at 8 µM FtsZ_AT_, n=3). Surprisingly, we did not observe polymerization of wildtype FtsZ_1_, even at high protein concentrations either in light scattering (Figure 3B, red line), TEM (Figure 3C, center panel), or GTP hydrolysis assays (Figure 3D, red line).

In light of the dependence of FtsZ_1_ on FtsZ_AT_ for mid-cell localization, we next sought to determine if FtsZ_AT_ and FtsZ_1_ can form co-polymers. To conduct these experiments, FtsZ_1_-L71W and FtsZ_AT_-L72W were purified to enable monitoring of protein polymerization using tryptophan fluorescence. The leucine to tryptophan mutation introduces a tryptophan on the surface of FtsZ that increases in fluorescence when it is buried in the subunit interface upon polymerization [40]. While wildtype FtsZ_AT_ (with no tryptophan) does not change in fluorescence on addition of GTP (Figure 3E, solid blue line), FtsZ_AT_-L72W fluorescence increases rapidly after GTP addition reflecting polymerization (Figure 3E, dashed blue line). When wildtype FtsZ_AT_ is added to FtsZ_AT_-L72W, bringing the total FtsZ concentration to 8 µM, fluorescence again increases, but then drops back to baseline upon complete consumption of GTP by this high concentration of FtsZ (Figure 3E, dotted blue line). Conversely, on its own or combined with wildtype FtsZ_1_, FtsZ_1_-L71W maintains a constant tryptophan fluorescence level before and after addition of GTP, consistent with our conclusion that it does not polymerize on its own (Figure 3E, red lines). Remarkably, tryptophan fluorescence increases when FtsZ_1_-L71W and FtsZ_AT_ are mixed, indicating that the FtsZ_1_-L71W is incorporated into polymers in the presence of FtsZ_AT_ (Figure 3C, purple dashed line). When FtsZ_AT_-L72W is mixed with FtsZ_1_, fluorescence increases above the level observed for FtsZ_AT_-L72W alone and drops to baseline faster than FtsZ_AT_-L72W on its own, again indicating co-polymerization. Finally, equimolar concentrations of FtsZ_AT_ alone or mixtures of FtsZ_AT_ and FtsZ_1_ exhibit similar rates of GTP hydrolysis (Figure 3D) and form qualitatively similar polymers by TEM (Figure 3C, right panel). Together, these observations indicate that FtsZ_1_ cannot polymerize independently, but that FtsZ_AT_ and FtsZ_1_ form co-polymers with similar structure and GTP hydrolysis rates as FtsZ_AT_ polymers.

Though multiple FtsZs are present in a number of bacterial and chloroplast lineages, their co-assembly properties have only begun to be characterized. In contrast to our observations, each of the FtsZs of *M. gryphiswaldense* was able to independently polymerize *in vitro*, but they also appeared to directly interact, perhaps reflecting an ability to co-polymerize [39]. Chloroplast FtsZs from *Arabadopsis thaliana* are also able to co-polymerize and, at least under some conditions, to independently polymerize [41]. Conversely, one of the FtsZs from tobacco chloroplasts cannot polymerize on its own but promotes polymerization of its partner homolog [42]. Finally, the FtsZ pair from the chloroplasts of representative green and red algae co-polymerize into polymers with altered assembly dynamics from either homopolymer [43]. It is likely that in each of these cases, the assembly or co-assembly properties of the duplicated FtsZs have evolved to suit a niche regulatory function. We hypothesize the FtsZ_1_ from *A. tumefaciens* has low affinity for itself, but higher affinity for FtsZ_AT_, limiting its homopolymerization but allowing for co-polymerization both *in vitro* and in cells. Since FtsZ_1_ cannot polymerize independently, FtsZ_AT_ must first polymerize at mid-cell after which FtsZ_1_ can be recruited by co-polymerization. The biological relevance of these biochemical and cell biological properties awaits further study.

### FtsZ_AT_ depletion results in tip splitting events

Once we identified FtsZ_AT_ as the primary homolog involved in cell division we next analyzed the growth phenotype during FtsZ_AT_ depletion more carefully. Compared to FtsZ_AT_ induced cells (Figure 4A, top), observation of cells during FtsZ_AT_ depletion by time-lapse microscope reveals remarkable changes in cell morphology (Figure 4A, bottom; Movie 1). Early during the depletion of FtsZ, an ectopic pole forms near mid-cell. We hypothesize that this occurs due to the ability of the remaining FtsZ to identify the mid-cell and recruit PG biosynthesis machinery to that site. Both the original growth pole and the ectopic pole are growth-active, resulting in the presence of multiple growth poles. These growth poles are unable to terminate cell elongation and ultimately most growth active poles are split, leading to the accumulation of many growth active poles (Figure 4A, bottom; Movie 1) and the rapid increase in cell area until the cell lyses.

**Figure 4.**
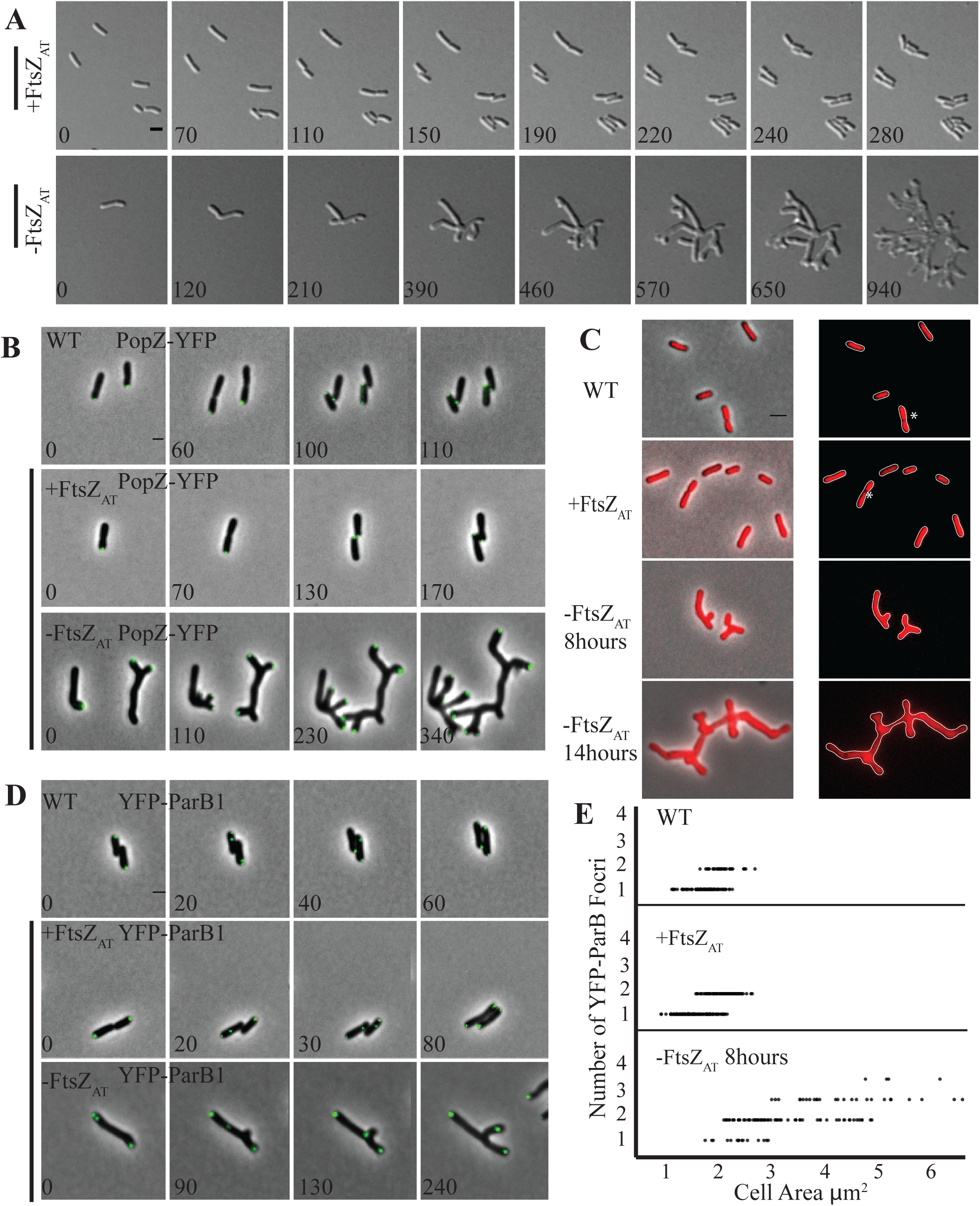
Characterization of genomic content during FtsZ_AT_ depletion. A) Timelapse microscopy in minutes demonstrates proper cell division and microcolony formation in +*ftsZ*_*AT*_ induced with IPTG (top panel). Timelapse during depletion of FtsZ_AT_ demonstrates branches forming from tip splitting events (bottom panel). B) Timelapse microscopy shows that PopZ-YFP maintains polar localization during elongation and dissociates moving to the mid-cell at division in WT and when *ftsZ*_*AT*_ is induced. PopZ-YFP becomes trapped at the growth poles during FtsZ_AT_ depletion. C) Sytox Orange labeled DNA is diffuse throughout young cells and separated nucleoids are seen in late divisional cells in WT and when FtsZ_AT_ is induced. These DNA free regions are marked with an asterisk. Nucleoids fail to form, as diffuse DNA labeling is observed during FtsZ_AT_ depletion at both 8 h and 14 h depletion timepoints. D) Timelapse microscopy demonstrates YFP-ParB1 form a single focus at the old pole in WT and when *ftsZ*_*AT*_ is induced. A second focus then translocates the cell length to the growth pole. Timelape microscopy during FtsZ_AT_ depletion reveals a third focus which translocates the cell towards an ectopic pole. E) Quantitation of YFP-ParB1 foci plotted against cell area. Number of foci increase as cell area increases. All scale bars are set to 2 µm.

The branched morphology observed during FtsZ_AT_ depletion is in stark contrast to FtsZ depletion observed in other organisms. In species like *E. coli* and *B. subtilis*, which utilize laterally localized peptidoglycan biosynthesis during elongation, depletion of FtsZ results in long, smooth filamentous cells. We hypothesize that the branching morphology of the *A. tumefaciens* FtsZ_AT_ depletion strain can be attributed to polar elongation. During the block in cell division, the growth pole continues to grow and presumably recruits additional peptidoglycan biosynthesis proteins. This could lead to an over-accumulation of elongasome proteins causing the pole to split into two poles. A similar branching pattern has been characterized during typical growth of *Streptomyces coelicolor* [44]. In this polar growing bacterium, the established elongasome splits, leaving a small portion of the elongasome behind as growth continues. With time, the subpolar elongasome accumulates in size and eventually forms a new growth pole. Although the polar growth molecular mechanisms are not conserved between *A. tumefaciens* and *S. coelicolor*, the fundamental principle of tip splitting as a consequence of polar growth appears to be shared.

### PopZ-YFP accumulates at growth poles in the absence of FtsZ_AT_

In WT *A. tumefaciens*, deletion of *popZ* has been shown to cause ectopic poles and cells devoid of DNA, demonstrating a role in coordinating cell division with chromosome segregation [45, 46]. We hypothesize that PopZ-dependent coordination of cell division likely involves FtsZ. In WT, PopZ-YFP localizes to the growing pole during elongation and is recruited to mid-cell just prior to cell separation (Figure 4B, top panel) [45-47]. When FtsZ_AT_ is expressed in the *ftsZ*_*AT*_ depletion strain, PopZ-YFP has a similar localization pattern as in WT cells (Figure 4B, middle panel). When FtsZ_AT_ is depleted, PopZ-YFP stays at the growth poles and as tip splitting events lead to the production of new growth poles, PopZ-YFP appears to be split and retained at all growth active poles (Figure 4B, bottom panel). These observations indicate that FtsZ is required for mobilizing PopZ from the growth pole to mid-cell. Remarkably, both FtsZ and FtsA are mislocalized in the absence of PopZ, leading to the establishment of asymmetric constrictions sites and a broad range of cell lengths [45]. Together, these data suggest that the presence of both PopZ and FtsZ are important for proper positioning and functioning of the divisome.

In addition to its function in maintaining proper cell division, *A. tumefaciens* PopZ is also required for chromosome segregation and tethers the centromere of at least one chromosome to the growth pole [46]. Thus, we examined the DNA content of cells depleted of FtsZ_AT_. In both WT cells and in conditions where *ftsZ*_*AT*_ is induced in the *ftsZ*_*AT*_ depletion strain, DNA labeled with Sytox orange is diffuse throughout most cells (Figure 4C, top two panels). In late divisional cells, true separation of nucleoids is observed indicating successful completion of chromosome segregation (Figure 4C, marked with an asterisk in the top two panels). In cells depleted of FtsZ_AT_ for both 8 and 14 h, DNA is diffuse throughout the elongated branches (Figure 4C, bottom two panels). The absence of distinct nucleoids may suggest that final stages of chromosome segregation are coordinated with cell separation as has been described for other bacteria including *E. coli* and *C. crescentus* [48].

To look more carefully at genomic content, we visualized YFP-ParB1, which serves as a proxy to track centromere partitioning in *A. tumefaciens* [46], in WT and *ftsZ*_*AT*_ depletion cells. In both WT cells and cells expressing FtsZ_AT_ in the *ftsZ*_*AT*_ depletion strain, a single YFP-ParB1 focus is present at the old pole in new cells generated by a recent cell division event (Figure 4D, top and middle panels). As the cells elongate, a second focus appears and translocates across the longitudinal axis to the growing pole (Figure 4D, top and middle panels). After 4 h of FtsZ_AT_pre-depletion, YFP-ParB1 foci can be seen at both poles, but when the cell fails to divide, a third focus of YFP-ParB1 appears and translocates along the longitudinal axis of the cell before taking a rapid turn toward a new ectopic pole formed from near mid-cell (Figure 4D, bottom panel). Next, we quantified the number of YFP-ParB1 foci relative to cell area (Figure 4E). In WT and FtsZ_AT_ expressing cells in the *ftsZ*_*AT*_ depletion strain, small cells have only a single focus of YFP-ParB1. This is followed by a transition period in which elongating cells accumulate a second focus of YFP-ParB1. Finally, the largest, pre-divisional cells have two YFP-ParB1 foci (Figure 4E). Cells depleted of FtsZ_AT_ for 8 h accumulate YFP-ParB1 foci as they increase in area (Figure 4E). Cells with an area larger than 3 μm^2^ all have at least 2 YFP-ParB1 foci, suggesting that chromosome replication is not blocked during FtsZ depletion. Furthermore, in larger cells additional YPF-ParB1 foci accumulate. These data suggest that cell division is not strictly required for the initiation of DNA replication in *A. tumefaciens*, although completion of chromosome segregation may be coordinated with cell division.

### Loss of septal PG synthesis results in altered total PG composition

Since polar growth appears to continue in the absence of FtsZ (Figure 4A, bottom panel), we used fluorescent-D-amino acids (FDAAs), to probe sites enriched in peptidoglycan synthesis [49] during depletion of FtsZ_AT_. In WT cells, FDAAs localize at a single pole in elongating cells and at mid-cell in pre-divisional cells (Figure 5A) [49]. As FtsZ_AT_ is depleted, FDAAs are targeted strictly to the poles, confirming that polar peptidoglycan synthesis is responsible for the observed increase in cell biomass after 8 h and 14 h of depletion (Figure 5A).

**Figure 5.**
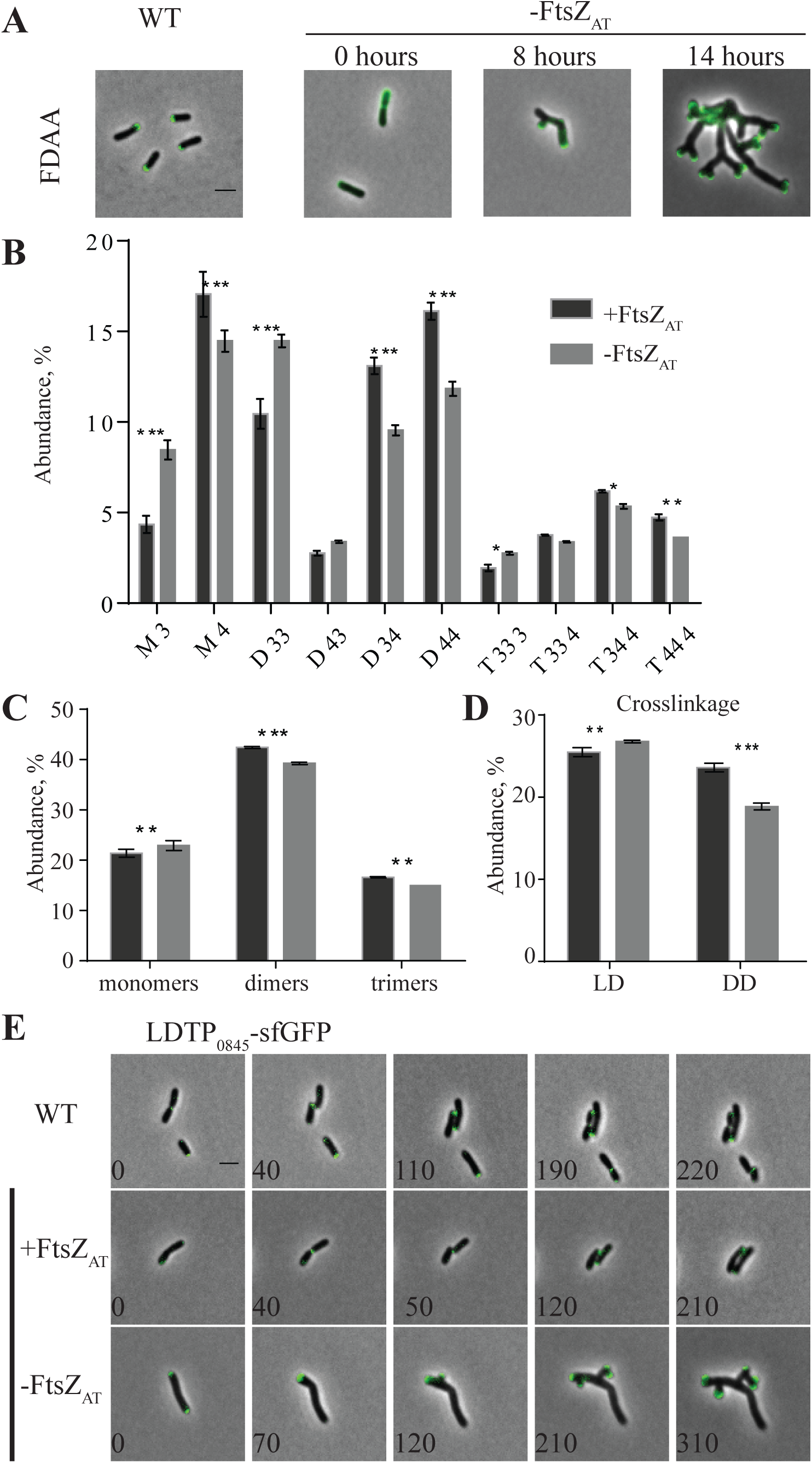
Characterization of polar peptidoglycan synthesis during FtsZ_AT_ depletion. A) FDAA labels active peptidoglycan synthesis at a single growing pole and septum in WT and cells depleted of FtsZ_AT_ for 0 h. As FtsZ_AT_ is depleted for 8 and 14 h, multiple growth poles are labeled, and septum labeling is lost. B) Quantitation of the major muropeptide peaks in *ftsZ*_*AT*_ depletion strain induced and depleted. C) Abundance of total monomers, dimers, and trimers in the muropeptide profile in *ftsZ*_*AT*_ depletion strain. D) Abundance of total LD and DD crosslinkage in *ftsZ*_*AT*_ depletion strain induced. For B,C,and D, data shown are the average abundance of each muropeptide and are taken from analysis of three independent biological samples of *ftsZ*_*AT*_ depletion strain induced (black bars) and depleted for 14 h (gray bars). Statistical was calculated by t-tests and is indicated with an asterisk (P-value <0.05 (*), <0.005 (**), <0.001 (***)). E) Timelapse microscopy of LDTP_0845_-sfGFP in WT and when *ftsZ*_*AT*_ is induced yields growth pole localization during elongation and mid-cell localization during septum formation. In cells depleted of FtsZ_AT_, localization is trapped at the growing poles. All scale bars are set to 2 µm.

Since cells depleted of FtsZ_AT_ fail to terminate polar growth and do not produce septal peptidoglycan, we hypothesized that the peptidoglycan composition may reveal chemical signatures of peptidoglycan derived from polar growth. Thus, we characterized the peptidoglycan composition of both WT cells and the *ftsZ*_*AT*_ depletion strain in both the presence and absence of IPTG using ultra-performance liquid chromatography (UPLC) [50]. The major muropeptides found in WT *A. tumefaciens* PG and their quantification are shown in (Figure 5-Figure Supplement 1) and include monomeric (M), dimeric (D), and trimeric (T) muropeptides. The muropeptide composition and abundance is similar between WT cells, WT cells grown in the presence of IPTG, and the *ftsZ*_*AT*_ depletion strain grown in the presence of IPTG such that FtsZ_AT_ is expressed (Figure 5-Figure Supplement 1). These findings suggest that there are no major changes in PG composition due to IPTG and that the presence of IPTG leads to complementation in the *ftsZ*_*AT*_ depletion strain. In contrast, when the *ftsZ*_*AT*_ depletion strain is grown in the absence of IPTG for 14 h, marked changes in muropeptide composition are observed (Figure 5B-D). While the overall abundance of monomeric, dimeric, and trimeric muropeptides are not dramatically impacted (Figure 5C), the abundance of specific muropeptides is modified. When FtsZ_AT_ is depleted, there is a significant increase in monomeric disaccharide tripeptide (M3) and a decrease in the abundance of the monomeric disaccharide tetrapeptide M4 (Figure 5B). This observation is consistent with the possibility that the absence of FtsZ_AT_ leads to an increase in LD-carboxypeptidase activity, which would remove the terminal peptide from M4, leading to both a reduction in the levels of M4 and an increase in the abundance in M3. Following FtsZ_AT_ depletion, the overall degree of muropeptide crosslinking decreases (Figure 5D). In particular, there is a marked decrease in DD-crosslinkages, which are formed by the DD-transpeptidase activity associated with penicillin-binding proteins (PBPs). The dominant dimeric muropeptide formed in the presence of FtsZ_AT_ is D44, which contains a DD-crosslink; in contrast, the dominant dimer formed in the absence of FtsZ_AT_ is D33, which contains an LD-crosslink (Figure 5B). These data suggest that the activity of LD-transpeptidases is increased and the activity of PBP-mediated DD-transpeptidases is decreased during FtsZ_AT_ depletion. The increased pool of M3 may provide additional acceptor substrate for LD-transpeptidases to increase the production of D33 relative to D44. In addition, increased LD-carboxypeptidase activity could contribute to increase further the levels of D33 using D34 as a substrate.

The *A. tumefaciens* genome contains 14 LD-transpeptidases, 7 of which are specific to Rhizobiales. The Rhizobiales-specific LD-transpeptidase encoded by *Atu_0845* (referred to here as LDTP_0845_) has been shown to localize to the growing pole in WT cells and has been hypothesized to contribute to polar growth [30]. This localization pattern was confirmed in both WT and FtsZ_AT_ induced cells (Figure 5E, top and middle panels). We find that LDTP_0845_ localizes at growth poles during depletion of FtsZ_AT_ (Figure 5E, bottom). This observation suggests that this LDTP_0845_ may contribute to changes in PG composition during FtsZ_AT_ depletion and supports a potential role for LD-transpeptidases in polar growth during elongation. The localization and function of putative periplasmic LD-carboxypeptidases in *A. tumefaciens* remain to be explored. Overall, these findings suggest that LD-carboxypeptidase and LD-transpeptidase activities are increased during FtsZ_AT_ depletion, indicating that these classes of enzymes may contribute to polar growth of *A. tumefaciens*.

### The C-terminal Conserved Peptide (CTP) of FtsZ_AT_ is required for proper termination of polar growth

To better understand the mechanism by which FtsZ_AT_ terminates polar growth, we constructed truncated proteins to analyze the function of the C-terminal conserved peptide (CTP) and the C-terminal linker (CTL) (Figure 1A). The CTP is a highly conserved domain which binds to proteins such as FtsA, that tether FtsZ to the membrane [37, 51, 52]. The CTL is an intrinsically disordered region of variable length found in FtsZ proteins, which functions in the regulation of PG biosynthesis and protofilament assembly [17, 53-55]. To probe the function of the FtsZ_AT_ CTP and CTL domains, we expressed FtsZ_AT_ΔCTP and FtsZ_AT_ΔCTL in both WT and FtsZ_AT_ depletion backgrounds.

In order to execute these experiments, we constructed a vector with an alternative “inducible” promoter system, which is compatible with the chromosomal IPTG depletion system. We modified pSRKKm [56] by replacing *lacI*^*q*^ with the gene encoding the cumate responsive repressor CymR [57, 58] and replacing the *lacO* operator sites with *cuO* operator sites (Figure 6-Figure Supplement 1A). This approach allows the same promoter to drive expression of both chromosomal full-length *ftsZ*_*AT*_ using IPTG and plasmid-encoded *ftsZ* variants using cumate. For simplicity, henceforth we referred to IPTG induction as mediated by P_lac_ and cumate induction as mediated by P_cym_. Expression of sfGFP from P_cym_ requires the presence of cumate (Figure 6-Figure Supplement 1B) and is comparable to expression of sfGFP from P_lac_ (Figure 6-Figure Supplement 1C). Although higher concentrations of cumate inhibit growth of WT *A. tumefaciens*, 0.01 mM cumate does not impair growth of WT cells (Figure 6-Figure Supplement 1D; left) and is sufficient to complement growth of the *ftsZ*_*AT*_ depletion strain in the absence of IPTG (Figure 6-Figure Supplement 1D; right).

**Figure 6.**
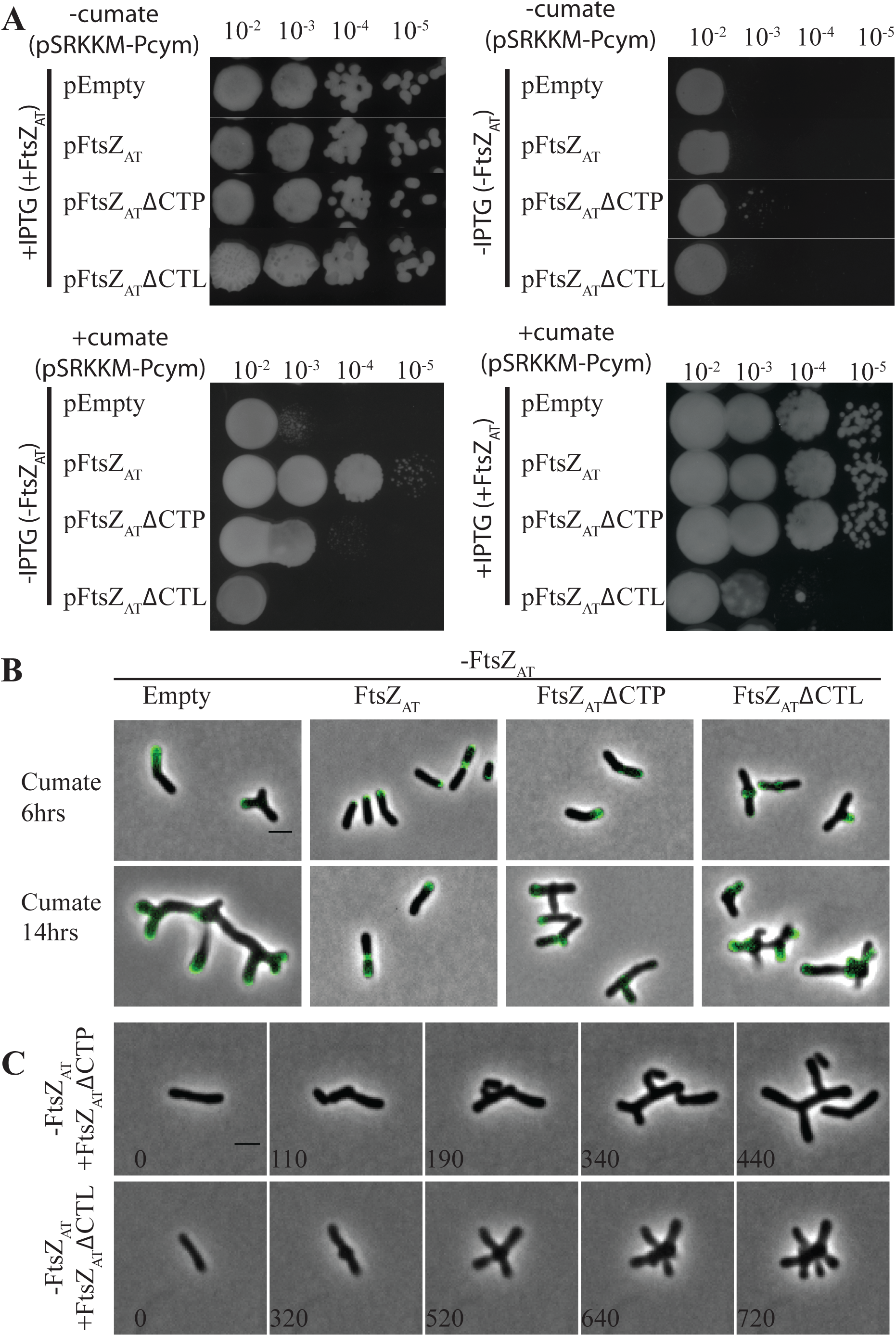
Functional analysis of FtsZ_AT_ΔCTL and FtsZ_AT_ΔCTP. A) Cell viability is measured by spotting serial dilutions of *ftsZ*_*AT*_ variants in the *ftsZ*_*AT*_ depletion background. When chromosomal *ftsZ*_*AT*_ is induced by IPTG and plasmid driven *ftsZ*_*AT*_ variants are uninduced, all strains have similar viability (top left). When both chromosomal *ftsZ*_*AT*_ is uninduced and plasmid driven *ftsZ*_*AT*_ variants are uninduced, all strains exhibit an equal decrease in viability (top right). When chromosomal *ftsZ*_*AT*_ is uninduced and plasmid driven *ftsZ*_*AT*_ variants are induced by cumate, FtsZ_AT_ expression rescues viability, FtsZ_AT_ΔCTP partially rescues, and FtsZ_AT_ΔCTL fails to rescue viability (bottom left). When both chromosomal *ftsZ*_*AT*_ is induced with IPTG and plasmid driven *ftsZ*_*AT*_ variants are induced by cumate, FtsZ_AT_ΔCTL expression reduces viability while other variants have no impact (bottom right). B) Representative images displaying morphology and FDAA labeling while chromosomal *ftsZ*_*AT*_ is uninduced and plasmid driven *ftsZ*_*AT*_ variants are induced for 6 and 14 h. C) Timelapse microscopy while chromosomal *ftsZAT* is uninduced and plasmid driven FtsZ_AT_ΔCTP is expressed reveal that polar growth fails to terminate and undergoes tip splitting, although septum formation and cell division also take place (top panel). Timelapse microscopy while chromosomal *ftsZ*_*AT*_ is uninduced and plasmid driven FtsZ_AT_ΔCTL is expressed shows termination of polar growth and new pole formation near mid-cell (bottom panel).

In the *ftsZ*_*AT*_ depletion strain, we introduced 4 vectors: an empty vector with P_cym_ (pEmpty), P_cym_- *ftsZ*_*AT*_ (pFtsZ_AT_), P_cym_-*ftsZ*_*AT*_*ΔCTP* (pFtsZΔCTP) or P_cym_-*ftsZ*_*AT*_*ΔCTL* (pFtsZΔCTL). When full-length *ftsZ*_*AT*_ is expressed from the chromosome, the viability of cells is not impacted by the presence of the P_cym_ vectors (Figure 6A, top left panel). In the absence of induction of *ftsZ* from the chromosome, the presence of the uninduced P_cym_ vectors, including pFtsZ_AT_, is not sufficient to rescue viability of the FtsZ-depleted cells (Figure 6A, top right panel); however, viability is significantly restored by expression of plasmid-encoded FtsZ_AT_ in the presence of cumate (Figure 6A, bottom left panel). Expression of plasmid-encoded FtsZ_AT_ΔCTP partially rescues the depletion of FtsZ_AT_ (Figure 6A, bottom left panel). In contrast, expression of plasmid-encoded FtsZ_AT_ΔCTL does not rescue the depletion of FtsZ_AT_ (Figure 6A, bottom left panel) and when both chromosomal full-length FtsZ_AT_ and FtsZ_AT_ΔCTL are expressed, viability is impaired, suggesting that FtsZ_AT_ΔCTL may have a dominant negative phenotype (Figure 6A, bottom right panel).

Next, we observed cell morphology of the *ftsZ*_*AT*_ depletion strain carrying each of the four vectors under conditions where the chromosomal FtsZ_AT_ is depleted and the plasmid-encoded FtsZ variants are expressed for 6 or 14 h (Figure 6B). The presence of pEmpty does not impact the FtsZ_AT_ depletion phenotype: branched cells with multiple growth active poles are observed (Figure 6B). Plasmid-encoded FtsZ_AT_ rescues the chromosomal FtsZ_AT_ depletion, resulting in the production of typical rod-shaped cells with PG biosynthesis occurring at a single pole or mid-cell (Figure 6B, middle left). The partial rescue of FtsZ_AT_ depletion in viability by expression of FtsZ_AT_ΔCTP was matched by a less severe defect in cell morphology (Figure 6B, middle right).

Although cells are branched, they are much shorter and have fewer branches than FtsZ_AT_ depletion. FDAA labeling reveals that the expression of FtsZ_AT_ΔCTP enables mid-cell labeling (Figure 6B, middle left), suggesting that PG is synthesized at mid-cell and that some cells may undergo division. Indeed, time-lapse microscopy of the FtsZ_AT_ depletion strain expressing only FtsZΔCTP reveals that the cells are capable of cell division events (Figure 6C, top panel). Remarkably, the sites of cell constriction and cell division are often asymmetric, giving rise to a cell population with a broad length distribution. Furthermore, polar growth is not terminated efficiently and both polar elongation and tip splitting events are evident. Together, these observations suggest that the FtsZ CTP contributes to proper termination of polar growth and divisome assembly. Expression of plasmid-encoded FtsZ_AT_ΔCTL in the absence of chromosome-encoded FtsZ_AT_ gives rise to a distinct cell morphology (Figure 6B, far right panel). After 6 hours of FtsZ_AT_ΔCTL expression, some cells contain mid-cell bulges. Remarkably, in these cells, FDAA labeling reveals that PG biosynthesis is occurring in the bulges and not at either pole. After 14 h, most cells have mid-cell swelling and multiple ectopic poles. Time-lapse microscopy reveals that polar growth is terminated and growth appears to be directed to mid-cell (Figure 6C, bottom panel, 320 min). When cell division fails, ectopic growth poles emerge from the mid-cell bulges (Figure 6C, bottom panel 520 min). The ectopic poles elongate, polar growth is terminated, and new ectopic growth poles are formed near the initial bulge site (Figure 6C, bottom panel). These observations suggest that the CTL of FtsZ_AT_ is required for proper cell division but is not required for the termination of polar growth.

### The CTL of FtsZ_AT_ is required for proper PG composition

The mid-cell bulges observed during FtsZΔCTL expression are reminiscent of those observed when FtsZ_CC_ΔCTL is expressed in *C. crescentus* [17]. In *C. crescentus*, the CTL was shown to be required for robust PG biosynthesis [17]. We therefore hypothesized that the altered PG composition observed during depletion of FtsZ_AT_ could be due to absence of the CTL. To test this hypothesis, we introduced plasmids containing no FtsZ (empty vector control, pEmpty), full-length FtsZ_AT_ (pFtsZ_AT_), or FtsZ_AT_ΔCTL (pFtsZ_AT_ΔCTL) into the *ftsZ* depletion strain. Each strain was grown under conditions in which expression of FtsZ_AT_ from the chromosomal copy is depleted and expression of the FtsZ variant (if present) from the plasmid is induced. PG was isolated from these strains following induction/depletion and analyzed. Induction of full-length FtsZ_AT_ from the plasmid yields lower levels of monomeric muropeptides compared to other strains, especially M3, and increased levels of dimeric and trimeric muropeptides, including D44 and T444 (Figure 7A-B). Overall the expression of full-length FtsZ_AT_ leads to an increased level of muropeptides with DD-crosslinks (Figure 7C). These observations indicate that expression of plasmid-encoded full-length FtsZ_AT_ compensates for the loss of FtsZ_AT_ from the chromosome. In contrast, the expression of FtsZ_AT_ΔCTL did not compensate for the loss of full-length FtsZ as the PG composition is more similar to the PG profile of FtsZ-depleted cells (Figure 7A-C). This observation suggests that the CTL of FtsZ_AT_ likely function in the regulation of proper PG biosynthesis at mid-cell.

**Figure 7.**
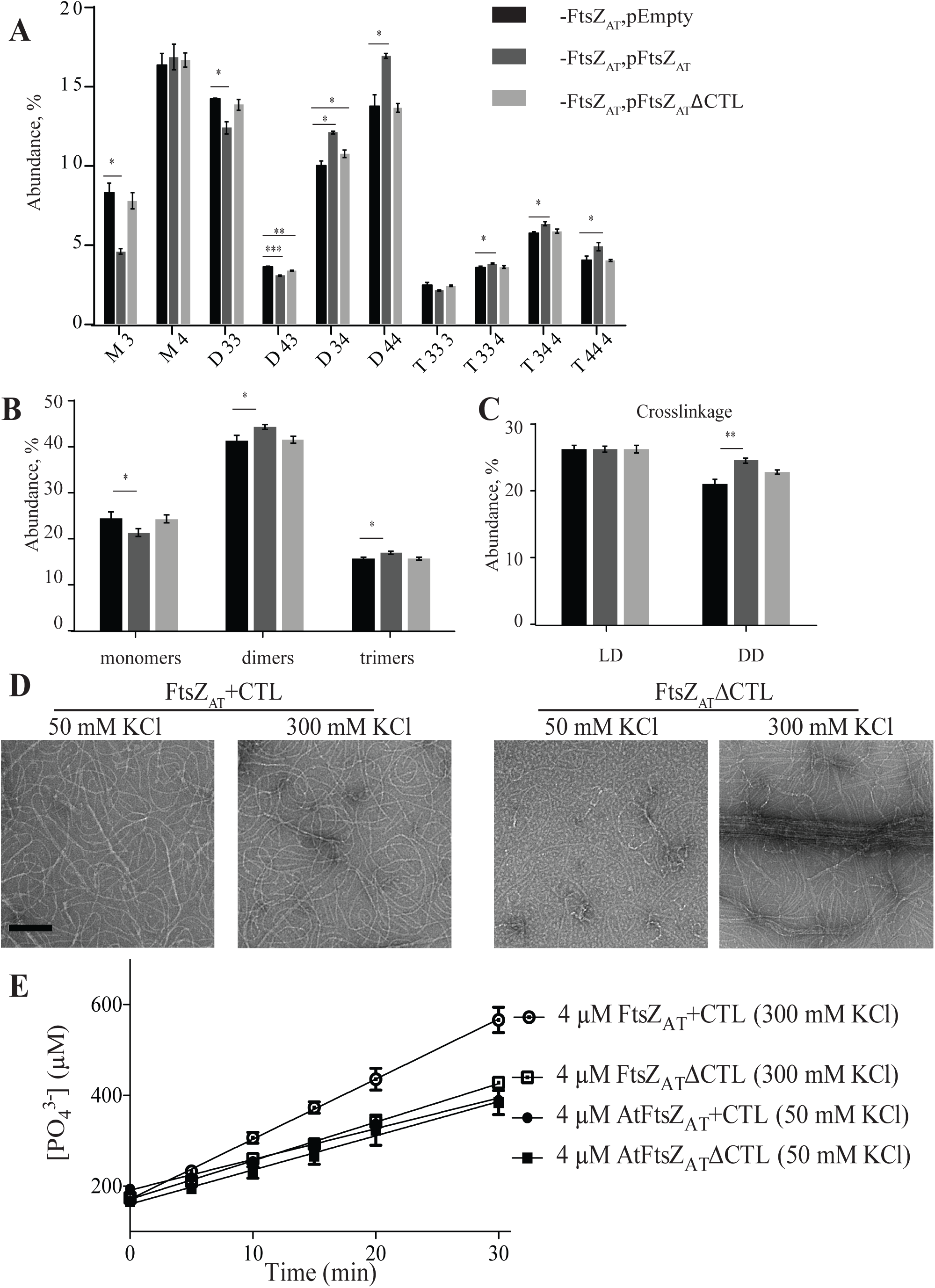
FtsZ_AT_ requires the CTL for robust PG biosynthesis and proper polymerization. A.) Quantitation of the major muropeptide peaks in *ftsZ*_*AT*_ depletion strain expressing an empty plasmid, full length FtsZ_AT_, or FtsZ_AT_ΔCTL. B) Abundance of total monomers, dimers, and trimers in the muropeptide profile. C) Abundance of total LD and DD crosslinkage. For A, B, and C, data shown are the average abundance of each muropeptide and are taken from analysis of three independent biological samples. Statistical significance is indicated with an asterisk. D.) Negative stain TEM of the 4 µM of the indicated protein in the presence of 50 or 300 mM KCl. FtsZ_AT_ΔCTL shows increased propensity to bundle at high salt. Scale bar is set at 100 nm. E.) Phosphate in solution over time in the presence of indicated proteins in solution with 50 or 300 mM KCl. The rate of GTP hydrolysis by FtsZ_AT_ΔCTL is reduced under high salt conditions that promote bundling.

### FtsZ CTL regulates protofilament assembly

Work in *C. crescentus* has shown that the FtsZ_CC_CTL directly regulates protofilament structure and dynamics [53]. To determine if the CTL of FtsZ_AT_ similarly regulates its assembly, we purified FtsZ_AT_ΔCTL and a control FtsZ_AT_+CTL protein containing the same restriction sites at the junctions with the GTPase domain and CTP as the ΔCTL construct, but with the CTL in place. FtsZ_AT_+CTL formed mostly single, gently curved protofilaments when visualized by TEM under all conditions tested (Figure 7D), similar to those observed for wildtype FtsZ_AT_ (Figure 3C). In contrast, under high salt conditions we observed extended bundles of FtsZ_AT_ΔCTL (Figure 7D). Furthermore, we saw a decreased rate of GTP hydrolysis by FtsZ_AT_ΔCTL under conditions that promote bundling (Figure 7D; 3.3 ± 0.2 GTP min^-1^ FtsZ^-1^ for FtsZ_AT_+CTL and 2.1 ± 0.1 GTP min^-1^ FtsZ^-1^ for FtsZ_AT_ΔCTL with 300 mM KCl, n =3). Together, these results suggest an important role for the CTL in limiting lateral interactions between protofilaments and promoting polymer turnover. These results in *A. tumefaciens* are consistent with effects of the CTL on polymer bundling reported in *C. crescentus* [17, 53]and *E. coli* [59]. Moreover, in light of our observations that FtsZ_AT_ΔCTL does not restore proper PG chemistry to FtsZ_AT_-depleted cells (Figure 7A,B), these data are in line with the growing body of evidence linking FtsZ dynamics and polymer superstructure to the regulation of PG biosynthesis.

### FtsA is required for cell division but not termination of polar growth

FtsA is an actin-like protein that associates with the membrane through an amphipathic helix and binds the FtsZ CTP to anchor FtsZ polymers to the membrane [51, 60]. In *C. crescentus*, recruitment of FtsA to mid-cell occurs well after the establishment of the FtsZ-ring and is dependent on the presence of FtsZ [13, 61]. In *A. tumefaciens*, FtsA-sfGFP is retained at the growth pole prior to appearing at mid-cell just before cell division [27, 30]. Here, we confirm that FtsA-sfGFP is observed as a focus at the growth pole until transitioning to a ring-like structure at mid-cell (Figure 8A, top panelIn fact, at some timepoints, both a polar focus and a mid-cell ring of FtsA are observed. Eventually, the polar focus disappears as the FtsA-sfGFP ring becomes more intense just prior to cell division. During the depletion of FtsZ_AT_, a focus of FtsA-sfGFP can be found at the growing pole, and at a newly formed ectopic pole near mid-cell (Figure 8A, bottom panel). FtsA-sfGFP remains associated with each growth pole, and as the poles undergo tip splitting events, each focus of FtsA-sfGFP is also split, resulting in the presence of FtsA-sfGFP in each of the 4 growth-active poles. These observations suggest that FtsZ_AT_ is required not only for proper mid-cell localization of FtsA to mid-cell prior to cell division but also contributes to release of FtsA-sfGFP from the growth pole.

**Figure 8.**
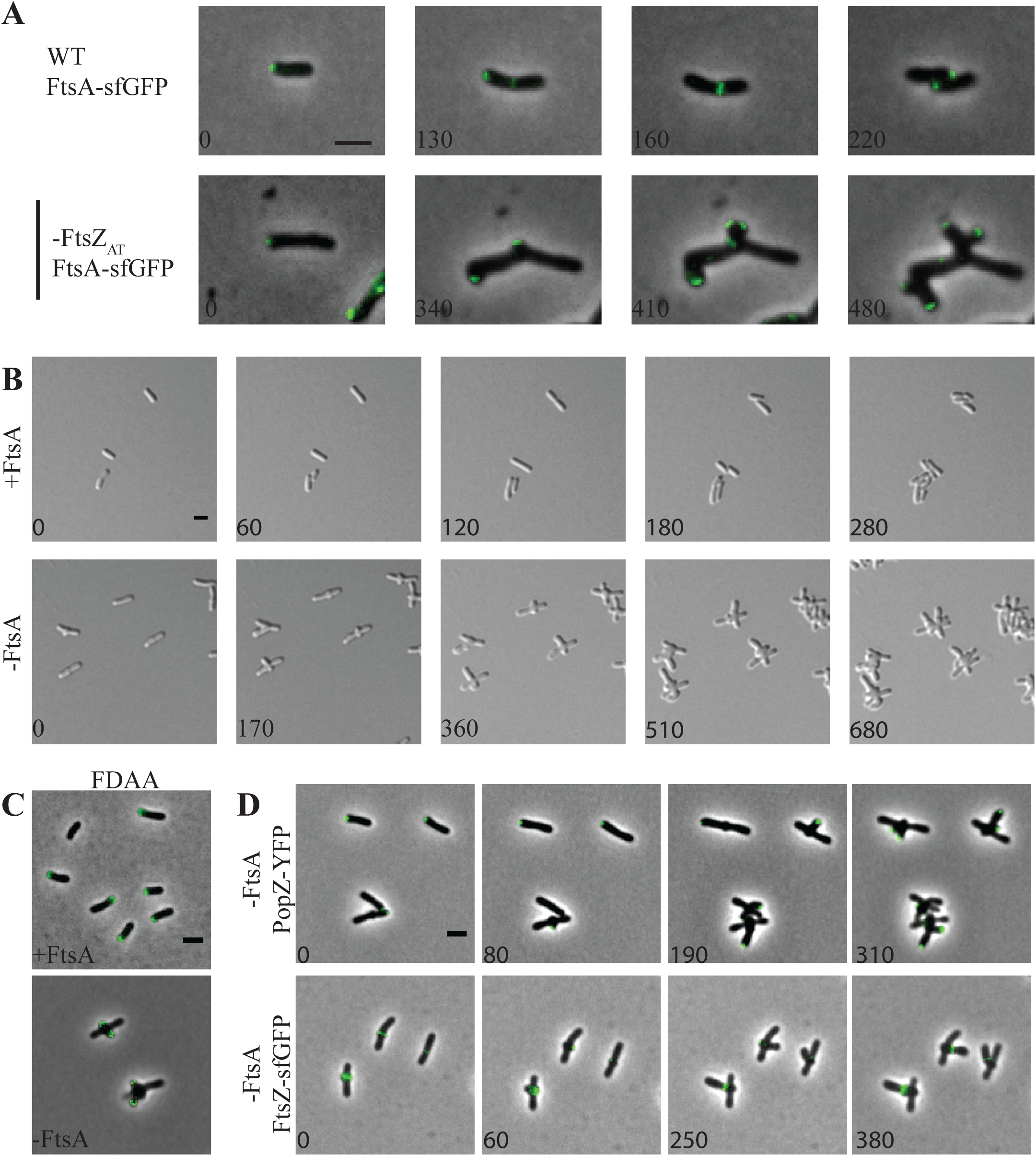
FtsA is not required for termination of polar growth. A) FtsA-sfGFP persists at growth poles and forms mid-cell rings which constrict in WT (top panel). FtsA-sfGFP becomes trapped at the growth poles and foci split as the growth poles split during FtsZ_AT_ depletion (bottom panel). B) Timelapse microscopy shows cells expressing FtsA grow and divide normally forming microcolonies (top panel). Cells depleted of FtsA terminate polar growth and form new growth poles near the mid-cell (bottom panel). C) FDAAs label a single growth pole when FtsA is present (top) and label multiple poles emerging from the mid-cell when FtsA is absent (bottom). D) Timelapse microscopy during FtsA depletion shows PopZ-YFP localizes to the growth poles and dissociates as growth is terminated. It then reappears at the new pole sites (top). During FtsA depletion, FtsZ-sfGFP forms rings marking the future sites of pole formation (bottom). All scale bars are set to 2 µm.

Since FtsA tethers FtsZ to the membrane and enables divisome assembly [37, 51, 52] in *E. coli*, we expected that the depletion of FtsA would phenocopy the depletion of FtsZ. Although a saturating transposon mutagenesis screen indicated that *ftsA* is not essential for *A. tumefaciens* cell survival [36], we were unable to construct a Δ*ftsA* mutant. Thus, we constructed a depletion strain in which expression of *ftsA* is controlled by P_lac_. Under conditions where FtsA is present in the *ftsA* depletion strain, cells maintain proper rod-shaped morphology, polar growth, and cell division occurs from constrictions formed near mid-cell (Figure 8B-C, top panels). In contrast, when FtsA is depleted, cells exhibit a marked change in morphology (Figure 8B, bottom panel; Movie 2). During the depletion of FtsA, rod-shaped cells initially elongate from a growth pole (Figure 8B, bottom panel, 0 min). Polar growth is terminated and growth is re-initiated from near mid-cell, typically resulting in the formation of two ectopic poles perpendicular to the original longitudinal axis of the cell (Figure 8B, bottom panel, 170 min). Cells depleted of FtsA continue multipolar growth (Figure 8B, bottom panel, 360 min), terminate growth from both poles and reinitiate growth from near mid-cell resulting in the formation of a new pair of ectopic growth poles (Figure 8B, bottom panel, 510 min). This pattern of multipolar growth, polar growth termination, and new branch formation is continued until cells eventually bulge at the mid-cell and lyse. Overall these observations indicate that the phenotypes caused by FtsZ and FtsA depletion are distinct from one another and suggest that only FtsZ is required for proper termination of polar growth.

To confirm that polar growth occurs and is terminated during FtsA depletion, cells were labeled with FDAAs (Figure 8C, bottom panel). Indeed, FDAAs label the tips of two poles, which are emerging from near mid-cell consistent with the re-initiation of polar growth. To further confirm that polar growth is terminated during FtsA depletion, we observed the localization of PopZ-YFP (Figure 8D, top panel). PopZ marks the growth poles [47] and becomes trapped at growth poles during depletion of FtsZ (Figure 4B). During FtsA depletion, PopZ-YFP is initially present at the growth pole (Figure 8D, top panel, 0 min). Next, PopZ-YFP disappears from the growth poles and reappears near mid-cell (Figure 8D, top panel, 80 min) indicating that polar growth is terminated. Throughout the FtsA depletion, PopZ-YFP continues to disappear from growth poles and reappears at the tips of newly emerging growth poles. Overall, these observations clearly indicate that FtsA is not necessary for termination of polar growth; however, FtsA has an essential function at a later stage of cell division since the cells fail to divide and are prone to lysis.

The ability of cells to target growth to near mid-cell during FtsA depletion suggests that FtsZ-rings may form, enabling the termination of polar growth. Indeed, FtsZ_AT_-sfGFP-rings form near mid-cell early during FtsA depletion (Figure 8D, bottom panel). FtsZ_AT_-sfGFP is briefly retained at new growth poles before reappearing to mark the site where a new growth pole will emerge. These observations are consistent with the finding the FtsA is retained at the growth pole longer than FtsZ [27, 62], and suggest that FtsA arrives at mid-cell after Z-ring assembly and the initiation of FtsZ-dependent cell wall biogenesis. The results observed here in *A. tumefaciens* are consistent with the observation that FtsA arrives to mid-cell after FtsZ and the onset of mid-cell cell wall biogenesis in *C. crescentus* [13, 61]. In both *A. tumefaciens* and *C. crescentus*, the late arrival of FtsA to the divisome suggests that other proteins contribute to proper tethering of FtsZ to the membrane. In *C. crescentus*, the FtsZ-binding protein, FzlC, functions as a membrane anchor early during the establishment of the divisome [31, 63]. A homolog of FzlC is readily found in the *A. tumefaciens* genome (Atu2824) and may contribute to the ability of FtsZ-rings to form in the absence of FtsA.

### Depletion of the downstream divisome component FtsW phenocopies depletion of FtsA

Having observed a distinct effect on cell morphology in the absence of *ftsA*, we wondered if the phenotype observed during *ftsA* depletion could be recapitulated in the absence of another late-arriving divisome protein. To test this hypothesis, we constructed a depletion strain of FtsW, which is recruited to mid-cell after FtsA in both *E. coli* and *C. crescentus* divisome assembly models [11, 13]. Depletion of FtsW results in a phenotype which is strikingly similar to the depletion of FtsA (Figure 9). When FtsW is induced normal growth is observed (Figure 9A, top panel). During FtsW depletion, polar growth is terminated, resulting in the establishment of growth-active poles from near mid-cell (Figure 9A, bottom panel; Movie 3). Multiple rounds of termination of polar growth followed by reinitiation of growth from near mid-cell occur until the mid-cell bulges and the cells ultimately lyse (Figure 9A, bottom panel). Labeling of growth active poles with FDAAs (Figure 9B) or by tracking PopZ-YFP localization (Figure 9C, top panel) confirmed that new branches which emerge from mid-cell are formed by polar growth.

**Figure 9.**
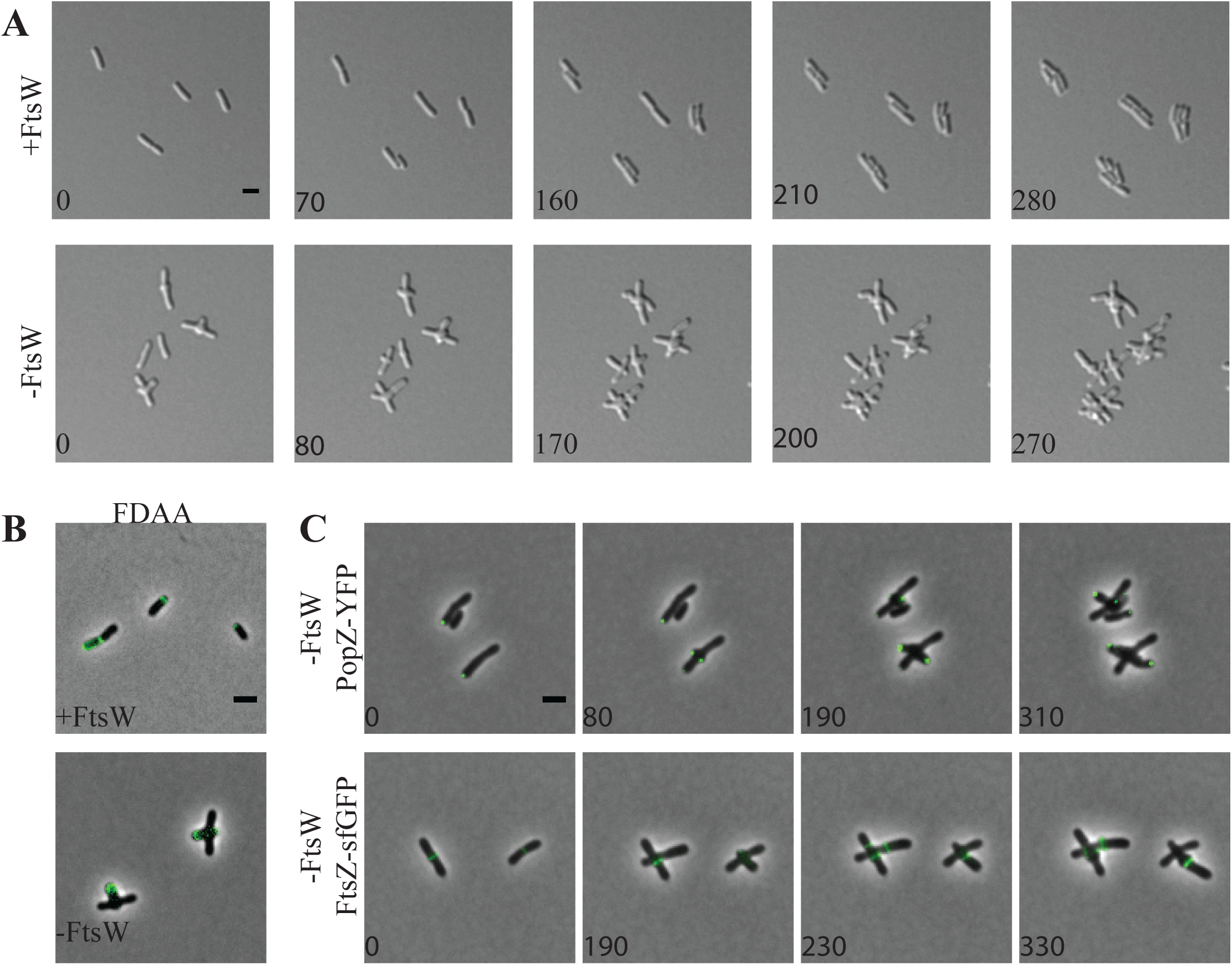
FtsW is not required for termination of polar growth. A) Timelapse microscopy shows cells expressing FtsW grow and divide normally forming microcolonies (top panel). Cells depleted of FtsW terminate polar growth and form new growth poles near the mid-cell (bottom panel). B) FDAA labels a single growth pole when FtsW is present (top) and labels multiple poles emerging from the mid-cell when FtsW is absent (bottom). C) Timelapse microscopy during FtsW depletion shows PopZ-YFP localizes to the growth poles and dissociates as growth is terminated. It then reappears at the new pole sites (top). During FtsW depletion, FtsZ-sfGFP forms rings marking the future sites of pole formation (bottom).

Finally, we confirmed that FtsZ-rings form during the depletion of FtsW and the presence of an FtsZ_AT_-sfGFP-ring typically marks the site where an ectopic growth pole will form (Figure 9C, bottom panel). Together, these observations suggest that FtsZ-rings are formed in the absence of FtsW, enabling the initiation of cell wall biogenesis. Given that FtsW drives septal PG biosynthesis, [64] these findings indicate that the cell wall biogenesis that occurs during depletion of FtsA or FtsW may require the elongation machinery, which typically functions in polar growth. Since the elongation machinery for *A. tumefaciens* remains to be identified, it is possible that there is considerable overlap between the machineries that contribute to polar and septal PG biosynthesis.

## Concluding Remarks

While many questions remain unanswered about the regulation of cell wall biogenesis in *A. tumefaciens*, our work sheds light on the transition from polar growth to mid-cell growth. We find that FtsZ_AT_, FtsA, and FtsW are required for constriction and cell separation, but FtsZ_AT_ is also required to terminate polar growth and initiate mid-cell peptidoglycan synthesis. How might the formation of an FtsZ_AT_-ring at mid-cell cause the termination of polar growth? We find that PopZ, and LDTP_0845_ become trapped at the growth poles during FtsZ depletion (Figure 5). It is possible that one or more of these proteins contributes to both polar peptidoglycan biosynthesis and mid-cell peptidoglycan synthesis and that the FtsZ-dependent targeting of these proteins (and likely others) to mid-cell triggers the termination of polar growth. While the mid-cell localization of PopZ is dependent on the presence of FtsZ_AT_ (Figure 4), FtsZ_AT_-ring stability and placement are impacted by the absence of PopZ [45]. Furthermore, deletion of *popZ* impairs termination of polar growth and results in cell division defects [45, 46, 65]. The apparent co-dependence of FtsZ and PopZ for localization may suggest that these proteins function together during the early stages of cell division, particularly the termination of polar growth and onset of mid-cell PG biosynthesis.

Overall, our results are consistent with a general model, which is highly conserved in bacteria, in which the establishment of a FtsZ-ring leads to the recruitment of many other cell division proteins to mid-cell [11], though many mechanistic questions remain. How is FtsZ_AT_ targeted to mid-cell? A variety of mechanisms that contribute to the proper placement of FtsZ at mid-cell have been described (for review see [66, 67]). The most well studied mechanisms of FtsZ positioning include negative regulation by the Min system and nucleoid occlusion. While genes encoding components of the Min system are readily identifiable in the *A. tumefaciens* genome, the deletion of *minCDE* has a minimal impact on placement of constriction sites and cell division efficiency [68]. Furthermore, FtsZ_AT_-GFP rings form over DNA prior to nucleoid separation in *A. tumefaciens*. These observations indicate that additional regulatory mechanisms must contribute to proper division site selection in *A. tumefaciens*. Following the appearance of FtsZ at mid-cell, how is the FtsZ_AT_-ring stabilized? In *E. coli*, the FtsZ-ring is stabilized by interactions with FtsA and ZipA, which tether FtsZ filaments to the membrane [51, 52, 69, 70]. In *A. tumefaciens*, FtsZ_AT_ appears at mid-cell well before FtsA [30] and we observe that FtsZ_AT_ rings form even when FtsA is depleted (Figure 7C, bottom panel). Furthermore, the position of FtsZ_AT_-GFP rings marks the site of ectopic pole formation. These observations suggest the FtsZ_AT_ is stabilized, at least early during cell division by other proteins. While there are no obvious ZipA homologs encoded in the *A. tumefaciens* genome, a homolog of FzlC, which functions to stabilize FtsZ in *C. crescentus* [31, 63], is encoded in the genome.

The observation that FtsZ is necessary for the initiation of mid-cell PG biosynthesis suggests that FtsZ is necessary for recruitment of PG biosynthesis enzymes to mid-cell. Septal PG biosynthesis is likely mediated by FtsW, a putative PG glycosyltransferase [71-73], and PBP3 (FtsI), a PG DD-transpeptidase [74]. In *A. tumefaciens*, depletion of FtsW does not cause a complete block of PG synthesis at mid-cell (Figure 8). This observation suggests that mid-cell PG biosynthesis is mediated by other cell wall biogenesis enzymes while the activity of FtsW contributes to later stages of cell division, consistent with the inability of cells to form constrictions and separate in the absence of FtsW. These observations may indicate that the initial PG biosynthesis at mid-cell comprises the final stage of cell elongation, consistent with descriptions of FtsZ-dependent mid-cell elongation in *C. crescentus* [16]. The observation that growth-active, ectopic poles emerge from near mid-cell during FtsW depletion (Figure 8B) provides evidence in support of this possibility. Thus, FtsZ-dependent PG biosynthesis may contribute to both elongation and cell division in *A. tumefaciens*. For a polar growing bacterium, it is tempting to speculate that the retention of PG biosynthesis machinery dedicated to elongation at the site of cell division may prime the newly formed poles to become growth active following cell separation.

## Materials and Methods

### Bacterial strains and culture conditions

All bacterial strains and plasmids used are listed in Table S4.1. *A. tumefaciens* strains were grown in ATGN minimal medium with .5% glucose [75] at 28°C. *E. coli* strains were grown in Luria-Bertani medium at 37°C. When indicated, kanamycin (KM) was used at 300 µg/ml for *A. tumefaciens*, 50 µg/ml for *E. coli* DH5α, and 25 µg/ml for *E. coli* S17-1 λ *pir*. Gentamicin was used when indicated at 200 µg/ml for *A. tumefaciens* and 20 µg/ml for *E. coli* DH5α. IPTG was added at a concentration of 1 mM when indicated. Cumate was added at a concentration of 0.1 mM when indicated.

### Construction of expression plasmids and strains

All strains and plasmids used are listed in Table S4.1, while primers used are listed in Table S4.2. For amplification of target genes, primer names indicate the primer orientation and added restriction sites. To construct expression vectors containing *ftsZ*_*AT*_*-sfgfp, ftsZ*_*1*_*-sfgfp, ftsZ*_*3*_*-sfgfp*, and *ldtp*_*0845*_*-sfgfp* the respective coding sequence was amplified from purified C58 genomic DNA using primers indicated in Table S4.2. The amplicons were digested overnight and ligated into cut pSRKKM-P_lac_-*sfgfp* using NEB T4 DNA ligase at 4°C overnight. The newly formed *sfgfp* fusion of each gene was excised from the plasmid by overnight digestion with NdeI and NheI. Fragments containing *ftsZ*_*AT*_*-sfgfp, ftsZ*_*1*_*-sfgfp, ftsZ*_*3*_*-sfgfp*, and *ldtp*_*0845*_*-sfgfp* were then ligated into cut pRV-MCS2 to give constitutive expression vectors containing the fusions. To construct the *popZ-yfp* expression vector, *popZ* along with the upstream promoter sequence were amplified from purified C58 genomic DNA, digested and ligated into pMR10.

To construct pSRKKM-P_cym_, a synthesized gBlock from IDT Integrated DNA Technologies was made containing the regulatory elements of the cumate system similar to previously described plasmid constructs [76, 77]. The P_cym_ promoter region is annotated in Table S4.2. The sequence encoding the cumate repressor was codon optimized for *A. tumefaciens* and placed under the control of the constitutive kanamycin promoter from pSRKKm-P_lac_-sf*gfp*. The synthesized gBlock was digested overnight with EcoRI and NdeI. The resulting fragment was then ligated into cut pSRKKm-P_lac_-sf*gfp* thereby replacing the original *lac* promoter and repressor with the cumate repressor and cumate regulated promoter.

Next, *yfp-parB* was excised from pSRKKM-P_lac_-*yfp-parB* [46] and ligated into pSRKKM-P_cym_ to create an expression vector compatible with the depletion strains. To create expression vectors for *ftsZ*_*AT*_, *ftsZ*_*AT*_*ΔCTP*, and *ftsZ*_*AT*_*ΔCTL* the respective target gene was amplified utilizing indicated primers, digested overnight with NdeI and BamHI and ligated into pSRKKM-P_cym_.

All expression vectors were verified by sequencing. All vectors were introduced into *A. tumefaciens* strains utilizing standard electroporation protocols [78] with the addition of IPTG in the media when introducing plasmids into in depletion backgrounds.

### Construction of deletion/depletion plasmids and strains

Vectors for gene deletion by allelic exchange were constructed using recommended methods for *A. tumefaciens* [78]. Briefly, 500 bp fragments upstream and downstream of the target gene were amplified using primer pairs P1/P2 and P3/P4 respectively. Amplicons were spliced together by SOEing using primer pair P1/P4. The amplicon was digested and ligated into pNTPS139. The deletion plasmids were introduced into *A. tumefaciens* by mating using an *E. coli* S17 conjugation strain to create KM resistant, sucrose sensitive primary exconjugants. Primary exconjugants were grown overnight in media with no selection. Secondary recombinants were screened by patching for sucrose resistance and KM sensitivity. Colony PCR with primers P5/P6 for the respective gene target was used to confirm deletion. PCR products from P5/P6 primer sets were sequenced to further confirm deletions.

For depletion strains, target genes (*ftsZ*_*AT*_, *ftsA,* and *ftsW*) were amplified, digested and ligated into either pUC18-mini-Tn*7*T-GM-P_lac_ or pUC18-mini-Tn*7*T-GM-P_lac_. The mini-Tn*7* vectors, along with the pTNS3 helper plasmid, were introduced into C58Δ*tetRA*::a-*att*Tn*7* as described previously [32]. Transformants were selected for gentamicin resistance and insertion of the target gene into the a-*att* site was verified by colony PCR using the tet forward and Tn7R109 primer. PCR products were sequenced to confirm insertion of the correct gene. Next, the target gene was deleted from the native locus as described above in the presence of 1 mM IPTG to drive expression of the target gene from the engineered site.

### Construction of plasmids for protein expression and purification

To construct pET21a FtsZ_AT_, *ftsZ*_*AT*_ was amplified from C58 genomic DNA with FtsZ_AT_ For NdeI and FtsZ_AT_ Rev EcoRI, digested with NdeI and EcoRI, and ligated into similarly digested pET21a. To construct pTB146 FtsZ_1_, *ftsZ*_*1*_ was amplified from C58 genomic DNA with FtsZ_1_ For SapI and FtsZ_1_ Rev BamHI, digested with SapI and BamHI, and ligated into similarly digested pTB146. Ligation products were transformed into NEB Turbo (New England Biolabs) and selected for ampicillin resistance. Insertions were verified by colony PCR and Sanger sequencing. Primers FtsZ_AT_-L72W and FtsZ1-L71W were used to mutagenize pET21a FtsZ_AT_ and pTB146 FtsZ_1_, respectively, using the Quikchange Multi Lightning Mutagenesis Kit (Agilent) and following the manufacturer’s protocol to generate pET21a FtsZ_AT_-L72W and pTB146 FtsZ_1_-L71W. Mutations in the targeted sites were verified by Sanger sequencing.

pET21c FtsZ_AT_+CTL and pET21c FtsZ_AT_ΔCTL were constructed in several steps. First, the GTPase domain of *ftsZ*_*AT*_ was amplified from C58 genomic DNA using FtsZ_AT_ For NdeI and FtsZ_AT_ GTPase Rev KpnI SacI, split into two aliquots, and digested with NdeI and KpnI or NdeI and SacI. The CTL region of *ftsZ*_*AT*_ was amplified from C58 genomic DNA using FtsZ_AT_ CTL For KpnI and FtsZ_AT_ CTL Rev SacI and digested with KpnI and SacI. For FtsZ_AT_+CTL, the GTPase domain amplicon (digested with NdeI and KpnI) was combined with the CTL amplicon (digested with KpnI and SacI) and together they were ligated into pXCFPN-1 [79] digested with NdeI and SacI. For FtsZ_AT_ΔCTL, the GTPase domain amplicon (digested with NdeI and SacI) was ligated into pXCFPN-1 digested with NdeI and SacI. Each was transformed into NEB Turbo, selected for spectinomycin resistance, and confirmed by colony PCR and Sanger sequencing. Next, the CTP was added to each of the above constructs by annealing oligos FtsZ_AT_ CTP + and FtsZ_AT_ CTP – (engineered with overhangs compatible with SacI and NheI ligation) and ligating into the above constructs digested with SacI and NheI. Each was transformed into NEB Turbo and confirmed as above to generate pX1 FtsZ_AT_+CTL and pX1 FtsZ_AT_ΔCTL. Finally, FtsZ_AT_+CTL and FtsZ_AT_ΔCTL were subcloned into pET21c by digestion of pX1 FtsZ_AT_+CTL and pX1 FtsZ_AT_ΔCTL with NdeI and NheI and ligating into similarly digested pET21c. Each was transformed into NEB Turbo, selected for ampicillin resistance, and confirmed by colony PCR and Sanger sequencing.

### DIC and phase contrast microscopy

Exponentially growing cells (OD_600_ = ∼0.6) were spotted on 1% agarose ATGN pads as previously described [80]. Microscopy was performed with an inverted Nikon Eclipse TiE with a QImaging Rolera em-c^2^ 1K EMCCD camera and Nikon Elements Imaging Software. For time-lapse microscopy, images were collected every ten minutes, unless otherwise stated.

### Fluorescence microscopy

Plasmid encoded FtsZ_AT_-sfGFP, FtsZ_1_-sfGFP, FtsZ_3_-sfGFP, and LDTP_0845_-sfGFP fusions were expressed from the P_van_ promoter, which provides constitutive low levels of expression (Figure 6-Figure Supplement 1C). Plasmid encoded FtsA-sfGFP and PopZ-YFP fusions were expressed from the native promoters. Expression of plasmid encoded YFP-ParB was induced by the presence of 0.1 mM cumate for 2 hours (h). Cells containing plasmids with fluorescent protein fusions were grown to exponential phase before imaging on agarose pads.

To visualize DNA, 1 ml of exponentially growing cells was treated with 1 µl of Sytox Orange for 5 minutes. Cells were collected by centrifugation and washed with PBS 2 times followed by a final resuspension in PBS. Cells were then imaged on agarose pads.

To visualize sites of active peptidoglycan synthesis 1 ml of exponentially growing cells was labeled with the fluorescent D-amino acid (FDAA), HCC amino-D-alanine (HADA), as previously described [49, 80].

### Cell viability and growth curve assays

For cell viability spot assays, exponentially growing cultures were diluted to OD_600_ = 0.1 and serially diluted in ATGN. 3 µl of each dilution was spotted onto ATGN and incubated at 28°C for 3 days before imaging. When appropriate ATGN plates contained KM 300 µg/ml, IPTG 1mM, and cumate 0.1 mM as indicated in figure legends. For growth curve analysis, exponentially growing cultures were diluted to OD_600_ = .05 in 200 µl of ATGN in 96-well plates. Plates were shaken for 1 minute before OD_600_ readings, which were taken every 10 minutes.

### Cell morphology and constriction rate analysis

Exponentially growing cells were imaged using phase contrast microscopy as described above. Cell length, area, and constrictions were detected using MicrobeJ software [81].

To calculate constriction rates, cells with detectable constrictions were tracked using time-lapse microscopy. The width of the cell constriction was measured at an initial time-point and the measurement was repeated after 10 minutes. The difference in constriction width was divided by the 10-minute time interval to give a constriction rate.

### Western blot analysis

For western blot analysis of FtsZ depletion, the *ftsZ* depletion strain was grown in 40 ml ATGN with 1 mM IPTG to exponential phase. 2 ml of culture was collected prior to depletion (time 0) by centrifugation at 10,000 x *g* for 3 minutes. The remaining culture was collected by centrifugation at 3500 x *g* for 10 minutes, and supernatants were discarded. Cells were washed in sterile water and pelleted again. To deplete FtsZ, the pellet was resuspended in fresh ATGN without IPTG and grown under standard culturing conditions. 2-ml samples were collected by centrifugation after 30, 45, 60, 120, and 240 minutes of depletion. OD_600_ was taken for each sample prior to centrifugation so that samples could be normalized to an OD_600_ equivalent to 0.68. The cell pellets were incubated with 100 μl of a master mix containing 1 ml of BugBuster protein extraction reagent (Novagen) and supplemented with 1 EDTA-free protease inhibitor cocktail (Sigma), 10 μl of lysonase (Novagen), 2,500 U/ml DNase I (Thermo Scientific), and 1 mM dithiothreitol (DTT) (Thermo Scientific) for 25 minutes with shaking at room temperature to lyse the cell pellets. The whole-cell lysates were clarified by centrifugation at 10,000 rpm for 15 min. A final concentration of 1 X Laemmli buffer was added to the cleared cell lysates. Samples were boiled at 100°C for 5 min prior to loading on a 4-15% Mini-PROTEAN TGX Precast Gel (Bio-Rad). The separated proteins were electroblotted onto polyvinylidene difluoride (PVDF) membranes (Bio-Rad) and blocked overnight in 5% nonfat dry milk powder solubilized in 1% TBST (Tris-buffered saline [TBS], 1% Tween 20). The blocked PVDF membranes were probed with *Escherichia coli* anti-FtsZ (1:3000) monoclonal antibody (gift from Joe Lutkenhaus) for 1.5 h in 5% milk-TBST, followed by incubation with anti-rabbit (1:5000) HRP (Pierce 31460) secondary antibody for 1 h in 5% milk-TBST. The secondary antibody was detected using the ECL Plus HRP substrate (Thermo Scientific Pierce).

For comparison of expression from P_van_, P_lac_, and P_cym_ promoters, strains were grown in 2 ml ATGN with 200 ug/mL KM to exponential phase. P_lac_ and P_cym_ were induced with 1 mM IPTG and 50 μM cumate, respectively for 4 h. Cell pellets were lysed as described above and clarified whole-cell lysates were boiled with 1 X Laemmli buffer for 5 min prior to loading on 4-15% Mini-PROTEAN TGX Precast Gel (Bio-Rad). The separated proteins were electroblotted onto PVDF membranes (Bio-Rad), blocked as described above, and probed with anti-GFP (1:3,000) monoclonal antibody (Thermo Scientific Pierce) for 1 h in 5% milk-TBST, followed by incubation with a donkey anti-mouse (1:300) horseradish peroxidase-conjugated secondary antibody (Thermo Scientific Pierce) for 1 h in 5% milk-TBST. The secondary antibody was detected using the ECL Plus HRP substrate (Thermo Scientific Pierce).

### Protein expression and purification

FtsZ_AT_, FtsZ_AT_-L72W, FtsZ_AT_+CTL (FtsZ_AT_ with restriction sites flanking the CTL), and FtsZ_AT_ΔCTL (FtsZ_AT_ with the CTL deleted, containing the same restriction sites at the GTPase-CTC junction as in FtsZ_AT_+CTL) were expressed and purified in untagged form. Each was produced from a pET21 expression vector (pEG1555 – FtsZ_AT_, pEG1556 - FtsZ_AT_-L72W, pEG1444 - FtsZ_AT_+CTL, pEG1445 - FtsZ_AT_ΔCTL) in *Escherichia coli* Rosetta(DE3)pLysS induced at 37°C for 4 h with 0.5 mM IPTG after OD reached 0.8 to 1.0 OD at 600 nm. Cells were harvested by centrifugation at 6000 x g and resuspended in 30 mL FtsZ QA buffer (50 mM Tris-HCl pH 8, 50 mM KCl, 0.1 mM EDTA, 10% glycerol) per liter of culture. Resuspensions were snap frozen in liquid nitrogen and stored at −80°C until purification. To purify, resuspensions were thawed quickly and cells were lysed by incubation with 1 mg/mL lysozyme, 2.5 mM MgCl_2_, DNAse I, 2 mM PMSF, and a cOmplete mini EDTA-free protease inhibitor tablet (Roche) for 45 min to 1 h at room temperature followed by sonication. Lysates were cleared by centrifugation at 15000 x g for 30 min at 4°C and filtered through a 0.45 µm filter before anion exchange chromatography (HiTrap Q HP 5 mL, GE Life Sciences). Protein was eluted with a linear KCl gradient (FtsZ QA buffer with 50 to 500 mM KCl) and fractions containing FtsZ were verified by SDS-PAGE, pooled, and subjected to ammonium sulfate precipitation. Precipitates (at 17-20% ammonium sulfate saturation depending on the variant) were verified by SDS-PAGE, resuspended in HEK50G (50 mM HEPES-KOH pH 7.2, 0.1 mM EDTA, 50 mM KCL, 10% glycerol, 1 mM β-mercaptoethanol), and further purified by gel filtration (Superdex 200 10/300 GL, GE Life Sciences). Peak fractions were pooled, snap frozen in liquid nitrogen, and stored at −80°C.

FtsZ_1_ and FtsZ_1_-L71W were produced as His_6_-SUMO fusions and cleaved to yield untagged, scarless proteins. Each was produced from a pTB146 expression vector (pEG1535 - FtsZ_1_, pEG1542 - FtsZ_1_-L71W) in *E. coli* Rosetta (DE3)pLysS as described above. Cells were harvested by centrifugation as above, resuspended in HK300G (50 mM HEPES-KOH pH7.2, 300 mM KCl, 10% glycerol) with 20 mM imidazole, snap frozen in liquid nitrogen, and stored at −80°C until purification. To purify, resuspensions were thawed quickly and cells were lysed by incubation with 1 mg/mL lysozyme, 2.5 mM MgCl_2_, and DNAse I for 45 min at room temperature followed by sonication. Lysate was cleared and filtered as described above. Protein was isolated by Ni^2+^ affinity chromatography (HisTrap FF 1 mL, GE Life Sciences) and eluted in HK300G with 300 mM imidazole. Fractions containing His_6_-SUMO fusions were verified by SDS-PAGE, combined with Ulp1 Sumo protease at a 1:100 (protease:FtsZ) molar ratio, and cleaved by incubation at 30°C for 3.5 h. Cleaved FtsZ_1_ or FtsZ_1_L71W was purified away from His_6_-SUMO by gel filtration (Superdex 200 10/300 GL, GE Life Sciences) in HEK50G. Peak fractions were pooled, snap frozen in liquid nitrogen, and stored at −80°C.

### Polymerization kinetics assays

A Fluoromax-3 spectrofluorometer (Jobin Yvon, Inc) was used to monitor FtsZ polymerization by right-angle light scattering and tryptophan fluorescence. FtsZ_1_ and/or FtsZ_AT_ (wild-type or L71W/L72W mutants, as indicated in figures and text) was polymerized in HEK50 (50 mM HEPES-KOH pH 7.2, 50 mM KCl, 0.1 mM EDTA) with 2.5 mM MgCl_2_. 2 mM GTP was used to induce polymerization for light scattering and 50 µM GTP was used to induce polymerization for tryptophan fluorescence (GTP is fluorescent at the wavelengths used, so low concentrations must be used). GTP was added after baseline light scatter or fluorescence was established. For light scattering, samples were excited at 350 nm and scatter was detected at 350 nm with slits set to 2 nm. For tryptophan fluorescence, samples were excited at 295 nm and emission was detected at 344 nm, with 2 nm slits.

### GTPase assay

FtsZ_1_ and/or FtsZ_AT_ was polymerized in HEK50 with 2.5 mM MgCl_2_ and 2 mM GTP. FtsZ_AT_+CTL or FtsZ_AT_ΔCTL was polymerized in HEK50 or HEK300 (same as HEK50 but with 300 mM KCl) as indicated, with 10 mM MgCl_2_ and 2 mM GTP. GTP was added at time 0. Reaction was stopped at 5, 10, 15, 20, and 30 minutes with quench buffer (50 mM HEPES-KOH pH 7.2, 21.3 mM EDTA, 50 mM KCl). Inorganic phosphate in solution (liberated by GTP hydrolysis) over time was measured using SensoLyte MG Phosphate Assay Kit Colorimetric (AnaSpec, Inc, Fremont, California).

### Negative stain transmission electron microscopy (TEM)

FtsZ_1_ and/or FtsZ_AT_ were polymerized in HEK50 with 2.5 mM MgCl_2_ and 2 mM GTP. 4 µM FtsZ_AT_+CTL or FtsZ_AT_ΔCTL were polymerized in HEK50 or HEK300 as indicated with 10 mM MgCl_2_ and 2 mM GTP. After a 15-minute incubation at room temperature, samples were applied to carbon-coated glow-discharged grids with 0.75% uranyl formate staining as previously described [17, 82]. TEM samples were imaged using a Philips/FEI BioTwin CM120 TEM equipped with an AMT XR80 8 megapixel CCD camera (AMT Imaging, USA).

### Peptidoglycan composition analysis

Six cultures of WT and *ftsZ* depletion cells were grown in 10 ml of ATGN with IPTG to exponential phase. The 10 ml cell cultures were added to 40 ml of fresh media. The 50 ml cultures were grown to exponential phase and pelleted by centrifugation at 4000 x *g* for 10 minutes. Cell pellets were washed three times with ATGN by centrifugation and resuspension to remove IPTG. After the final wash 3 cell pellets were resuspended in 50 ml ATGN and the remaining 3 pellets were resuspended in 50 ml ATGN with 1 mM IPTG. Each culture was grown for 14 h. The optical densities of the cells were monitored to ensure the optical density of the cultures never went above OD_600_ = 0.7 to avoid changes to peptidoglycan content due to stationary phase. If necessary, fresh medium was added to dilute the cultures to maintain exponential growth. After 14 h of growth, 50 ml of the exponential cultures were collected and pelleted by centrifugation at 4000 x *g* for 20 minutes. Cell pellets were resuspended in 1mL of ATGN and 2 mL of 6% SDS and stirred with magnets while boiling for 4 h. After 4 h, samples were removed from heat but continued to stir overnight. Samples were then shipped to Dr. Felipe Cava’s laboratory for purification and analysis.

Upon arrival, cells were boiled and simultaneously stirred by magnets for 2 h. After 2 h, boiling was stopped and samples were stirred overnight. Peptidoglycan was pelleted by centrifugation for 13 min at 60000 rpm (TLA100.3 Beckman rotor, Optima Max-TL ultracentrifuge; Beckman), and the pellets were washed 3 to 4 times by repeated cycles of centrifugation and resuspension in water. The pellet from the final wash was resuspended in 50 µl of 50 mM sodium phosphate buffer, pH 4.9, and digested overnight with 100 µg/ml of muramidase at 37°C. Muramidase digestion was stopped by boiling for 4 min. Coagulated protein was removed by centrifugation for 15 min at 15000 rpm in a desktop microcentrifuge. The muropeptides were mixed with 15 µl 0.5 M sodium borate and subjected to reduction of muramic acid residues into muramitol by sodium borohydride (10 mg/ml final concentration, 20 min at room temperature) treatment. Samples were adjusted to pH 3 to 4 with orthophosphoric acid and filtered (0.2-µm filters).

Muropeptides were analyzed on a Waters UPLC system equipped with an ACQUITY UPLC BEH C18 Column, 130 Å, 1.7 µm, 2.1 mm × 150 mm (Waters) and a dual wavelength absorbance detector. Elution of muropeptides was detected at 204 nm. Muropeptides were separated at 45°C using a linear gradient from buffer A [formic acid 0.1% (v/v)] to buffer B [formic acid 0.1% (v/v), acetonitrile 20% (v/v)] in a 12 min run with a 0.250 ml/min flow. Peptidoglycan compositional analysis on triplicate samples was completed on two separate occasions.

## ACKNOWLEDGEMENTS

We thank members of the Brown lab for helpful discussions and critical reading of this manuscript.

---

**Figure 1-Figure Supplement 1.**
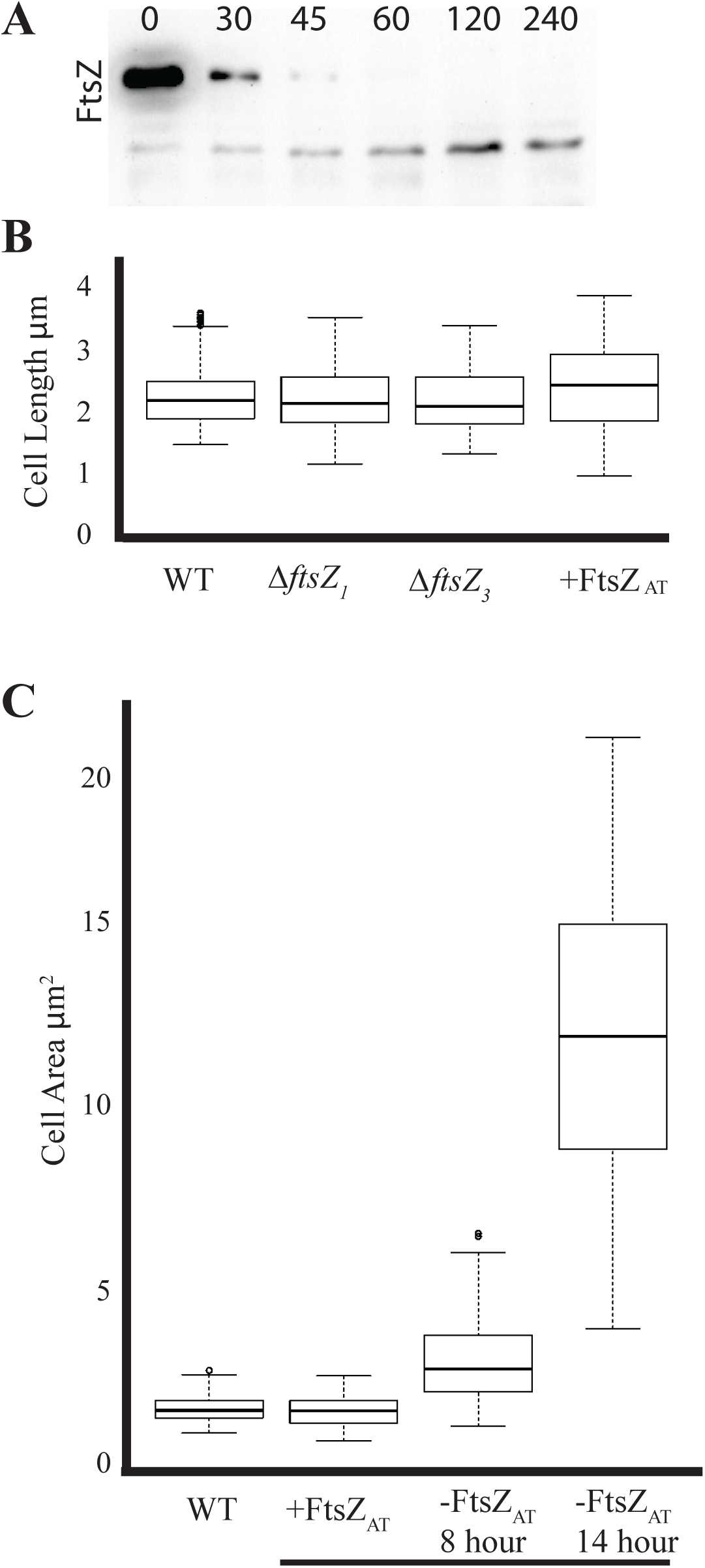
Cell growth and morphology of *ftsZ* mutants. A.) Western blot analysis showing the depletion of FtZ_AT_ after the removal of inducer. B) Cell lengths of WT, Δ*ftsZ*_*1*_, Δ*ftsZ*_*3*_, and induced *ftsZ*_*AT*_. are indistinguishable. C) Cell area in WT and induced *ftsZ*_*AT*_are the same while cells depleted of FtsZ_AT_ for 8 and 14 hours accumulate cell area.

**Movie 1. Growth and morphological changes during FtsZ_AT_ depletion.** Cells were washed to remove inducer and grown in liquid ATGN for 4 hours before spotting on a ATGN pad. Images were acquired every ten minutes and movie is played at 10 frames per second for a total of 145 frames.

**Figure 5-Figure Supplement 1.**
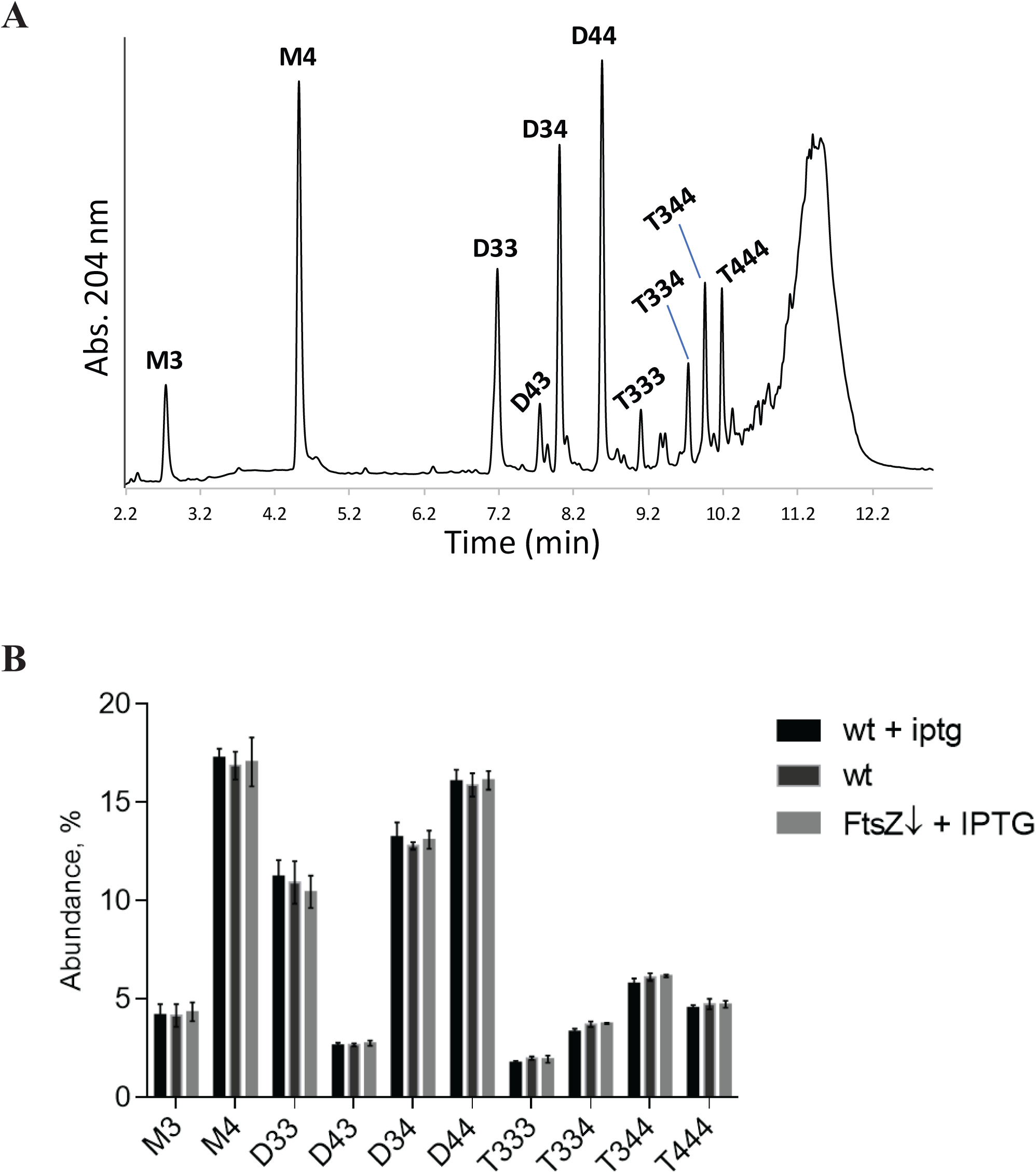
Peptidoglycan analysis of control strains. A) UPLC spectra of muropeptides derived from WT cells. Major muropeptides are labeled. M= monomers, D= dimers, T= trimers. Numbers indicate the length of the muropeptide stems and the position of crosslink in dimers and trimers. B) Quantitation of the major muropeptide peaks in WT with IPTG (black), WT without IPTG (black with gray outline), and *ftsZ*_*AT*_ depletion strain induced with IPTG (Gray). Data shown is the average abundance of each muropeptide and is taken from analysis of three independent biological samples. Statistical significance is indicated with an asterisk.

**Figure 6-Figure Supplement 1.**
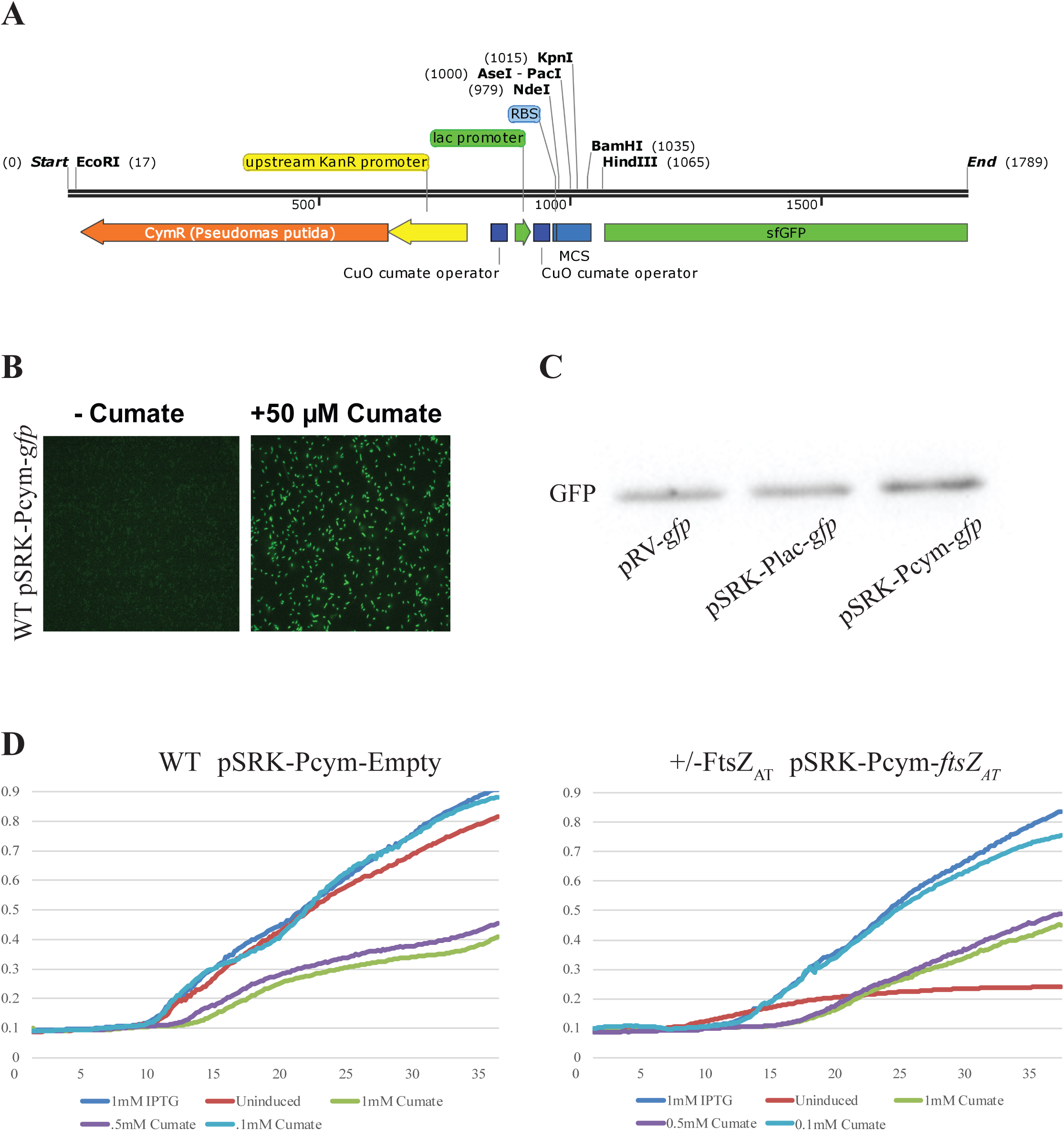
Validation of a cumate inducible vector in *A. tumefaciens.* A) Sequence schematic of the cumate operon modified for use is pSRKKM-*sfGFP*. Regions are color coded to match sequence found in Table S2. B) Representative image of WT cells harboring pSRKKM-Pcym-sfGFP uninduced (left) and induced (right). C) Western blot analysis comparing expression levels of sfGFP expressed from pRV, pSRKKM-Plac, and pSRKKM-Pcym. D) Growth curve analysis of WT cells harboring psRKKM-Pcym-empty induced with different levels of cumate (left). pSRKKM-Pcym-*ftsZ*_*AT*_ rescues chromosomal FtsZ_AT_ depletion with 0.01mM cumate (right).

**Movie 2. Growth and morphological changes during FtsA depletion.** Cells were washed to remove inducer and grown in liquid ATGN for 2 hours before spotting on a ATGN pad. Images were acquired every ten minutes and movie is played at 10 frames per second for a total of 97 frames.

**Movie 3. Growth and morphological changes during FtsW depletion.** Cells were washed to remove inducer and grown in liquid ATGN for 4 hours before spotting on a ATGN pad. Images were acquired every ten minutes and movie is played at 10 frames per second for a total of 85 frames.

## References

1. den Blaauwen T, Andreu JM, Monasterio O. Bacterial cell division proteins as antibiotic targets. Bioorg Chem. 2014;55:27–38. Epub 2014/04/24. doi: 10.1016/j.bioorg.2014.03.007. PMID: 24755375.

2. Haeusser DP, Levin PA. The great divide: coordinating cell cycle events during bacterial growth and division. Curr Opin Microbiol. 2008;11(2):94–9. Epub 2008/04/09. doi: 10.1016/j.mib.2008.02.008. PMID: 18396093.

3. Wu LJ, Errington J. Coordination of cell division and chromosome segregation by a nucleoid occlusion protein in *Bacillus subtilis*. Cell. 2004;117(7):915–25. Epub 2004/06/24. doi: 10.1016/j.cell.2004.06.002. PMID: 15210112.

4. Bi EF, Lutkenhaus J. FtsZ ring structure associated with division in *Escherichia coli*. Nature. 1991;354(6349):161–4. Epub 1991/11/14. doi: 10.1038/354161a0. PMID: 1944597.

5. de Boer P, Crossley R, Rothfield L. The essential bacterial cell-division protein FtsZ is a GTPase. Nature. 1992;359(6392):254–6. Epub 1992/09/17. doi: 10.1038/359254a0. PMID: 1528268.

6. Li Z, Trimble MJ, Brun YV, Jensen GJ. The structure of FtsZ filaments in vivo suggests a force-generating role in cell division. EMBO J. 2007;26(22):4694–708. Epub 2007/10/20. doi: 10.1038/sj.emboj.7601895. PMID: 17948052.

7. Fu G, Huang T, Buss J, Coltharp C, Hensel Z, Xiao J. In vivo structure of the *E. coli* FtsZ-ring revealed by photoactivated localization microscopy (PALM). PLoS One. 2010;5(9):e12682. Epub 2010/09/22. doi: 10.1371/journal.pone.0012680. PMID: 20856929.

8. Holden SJ, Pengo T, Meibom KL, Fernandez Fernandez C, Collier J, Manley S. High throughput 3D super-resolution microscopy reveals *Caulobacter crescentus* in vivo Z-ring organization. Proc Natl Acad Sci U S A. 2014;111(12):4566–71. Epub 2014/03/13. doi: 10.1073/pnas.1313368111. PMID: 24616530.

9. Bisson-Filho AW, Hsu YP, Squyres GR, Kuru E, Wu F, Jukes C, et al. Treadmilling by FtsZ filaments drives peptidoglycan synthesis and bacterial cell division. Science. 2017;355(6326):739–43. Epub 2017/02/18. doi: 10.1126/science.aak9973. PMID: 28209898.

10. Yang X, Lyu Z, Miguel A, McQuillen R, Huang KC, Xiao J. GTPase activity-coupled treadmilling of the bacterial tubulin FtsZ organizes septal cell wall synthesis. Science. 2017;355(6326):744–7. Epub 2017/02/18. doi: 10.1126/science.aak9995. PMID: 28209899.

11. Du S, Lutkenhaus J. Assembly and activation of the *Escherichia coli* divisome. Mol Microbiol. 2017;105(2):177–87. doi: doi:10.1111/mmi.13696.

12. Meier EL, Goley ED. Form and function of the bacterial cytokinetic ring. Curr Opin Cell Biol. 2014;26:19–27. Epub 2014/02/18. doi: 10.1016/j.ceb.2013.08.006. PMID: 24529242.

13. Goley ED, Yeh YC, Hong SH, Fero MJ, Abeliuk E, McAdams HH, et al. Assembly of the *Caulobacter* cell division machine. Mol Microbiol. 2011;80(6):1680–98. Epub 2011/05/06. doi: 10.1111/j.1365-2958.2011.07677.x. PMID: 21542856.

14. Erickson HP, Anderson DE, Osawa M. FtsZ in bacterial cytokinesis: cytoskeleton and force generator all in one. Microbiol Mol Biol Rev. 2010;74(4):504–28. Epub 2010/12/02. doi: 10.1128/mmbr.00021-10. PMID: 21119015.

15. Varma A, Young KD. FtsZ collaborates with penicillin binding proteins to generate bacterial cell shape in *Escherichia coli*. J Bacteriol. 2004;186(20):6768–74. Epub 2004/10/07. doi: 10.1128/jb.186.20.6768-6774.2004. PMID: 15466028.

16. Aaron M, Charbon G, Lam H, Schwarz H, Vollmer W, Jacobs-Wagner C. The tubulin homologue FtsZ contributes to cell elongation by guiding cell wall precursor synthesis in *Caulobacter crescentus*. Mol Microbiol. 2007;64(4):938–52. Epub 2007/05/16. doi: 10.1111/j.1365-2958.2007.05720.x. PMID: 17501919.

17. Sundararajan K, Miguel A, Desmarais SM, Meier EL, Huang KC, Goley ED. The bacterial tubulin FtsZ requires its intrinsically disordered linker to direct robust cell wall construction. Nat Commun. 2015;6:7281. doi: 10.1038/ncomms8281. PMID: 26099469.

18. Aarsman ME, Piette A, Fraipont C, Vinkenvleugel TM, Nguyen-Disteche M, den Blaauwen T. Maturation of the *Escherichia coli* divisome occurs in two steps. Mol Microbiol. 2005;55(6):1631–45. Epub 2005/03/09. doi: 10.1111/j.1365-2958.2005.04502.x. PMID: 15752189.

19. Gray AN, Egan AJ, Van’t Veer IL, Verheul J, Colavin A, Koumoutsi A, et al. Coordination of peptidoglycan synthesis and outer membrane constriction during *Escherichia coli* cell division. Elife. 2015;4. Epub 2015/05/08. doi: 10.7554/eLife.07118. PMID: 25951518.

20. Pini F, De Nisco NJ, Ferri L, Penterman J, Fioravanti A, Brilli M, et al. Cell cycle control by the master regulator CtrA in *Sinorhizobium meliloti*. PLoS Genetics. 2015;11(5):e1005232. doi: 10.1371/journal.pgen.1005232.

21. Howell M, Brown PJB. Building the bacterial cell wall at the pole. Current Opinion in Microbiology. 2016;34:53–9. doi: 10.1016/j.mib.2016.07.021.

22. Pini F, Frage B, Ferri L, De Nisco NJ, Mohapatra SS, Taddei L, et al. The DivJ, CbrA and PleC system controls DivK phosphorylation and symbiosis in *Sinorhizobium meliloti*. Mol Microbiol. 2013;90(1):54–71. Epub 2013/08/06. doi: 10.1111/mmi.12347. PMID: 23909720.

23. Bellefontaine AF, Pierreux CE, Mertens P, Vandenhaute J, Letesson JJ, De Bolle X. Plasticity of a transcriptional regulation network among alpha-proteobacteria is supported by the identification of CtrA targets in *Brucella abortus*. Mol Microbiol. 2002;43(4):945–60. Epub 2002/04/04. PMID: 11929544.

24. Cheng J, Sibley CD, Zaheer R, Finan TM. A *Sinorhizobium meliloti* minE mutant has an altered morphology and exhibits defects in legume symbiosis. Microbiology. 2007;153(Pt 2):375–87. Epub 2007/01/30. doi: 10.1099/mic.0.2006/001362-0. PMID: 17259609.

25. Latch JN, Margolin W. Generation of buds, swellings, and branches instead of filaments after blocking the cell cycle of *Rhizobium meliloti*. J Bacteriol. 1997;179(7):2373–81. Epub 1997/04/01. PMID: 9079925.

26. Fujiwara T, Fukui S. Effect OF D-alanine and mitomycin-c on cell morphology of *Agrobacterium tumefaciens*. Gen App Microbiol. 1974;20(6):345–9. doi: 10.2323/jgam.20.345.

27. Zupan JR, Cameron TA, Anderson-Furgeson J, Zambryski PC. Dynamic FtsA and FtsZ localization and outer membrane alterations during polar growth and cell division in *Agrobacterium tumefaciens*. Proc Natl Acad Sci U S A. 2013;110(22):9060–5. doi: 10.1073/pnas.1307241110. PMID: 23674672.

28. Figueroa-Cuilan WM, Brown PJB. Cell wall biogenesis during elongation and division in the plant pathogen *Agrobacterium tumefaciens*. Curr Top Microbiol Immunol. 2018. Epub 2018/05/29. doi: 10.1007/82_2018_92. PMID: 29808336.

29. Brown PJB, de Pedro MA, Kysela DT, Van der Henst C, Kim J, De Bolle X, et al. Polar growth in the Alphaproteobacterial order Rhizobiales. Proc Natl Acad Sci U S A. 2012;109(5):1697–701. doi: 10.1073/pnas.1114476109.

30. Cameron TA, Anderson-Furgeson J, Zupan JR, Zik JJ, Zambryski PC. Peptidoglycan synthesis machinery in *Agrobacterium tumefaciens* during unipolar growth and cell division. mBio. 2014;5(3). doi: 10.1128/mBio.01219-14. PMID: 24865559.

31. Goley ED, Dye NA, Werner JN, Gitai Z, Shapiro L. Imaging-based identification of a critical regulator of FtsZ protofilament curvature in *Caulobacter*. Mol Cell. 2010;39(6):975–87. Epub 2010/09/25. doi: 10.1016/j.molcel.2010.08.027. PMID: 20864042.

32. Figueroa-Cuilan W, Daniel JJ, Howell M, Sulaiman A, Brown PJ. Mini-Tn*7* Insertion in an artificial *att*Tn*7* site enables depletion of the essential master regulator CtrA in the phytopathogen *Agrobacterium tumefaciens*. Appl Environ Microbiol. 2016;82(16):5015–25. Epub 2016/06/12. doi: 10.1128/aem.01392-16. PMID: 27287320.

33. Ortiz C, Natale P, Cueto L, Vicente M. The keepers of the ring: regulators of FtsZ assembly. FEMS Microbiol Rev. 2016;40(1):57–67. Epub 2015/09/18. doi: 10.1093/femsre/fuv040. PMID: 26377318.

34. Wood DW, Setubal JC, Kaul R, Monks DE, Kitajima JP, Okura VK, et al. The genome of the natural genetic engineer *Agrobacterium tumefaciens* C58. Science. 2001;294(5550):2317–23. Epub 2001/12/18. doi: 10.1126/science.1066804. PMID: 11743193.

35. Goodner B, Hinkle G, Gattung S, Miller N, Blanchard M, Qurollo B, et al. Genome sequence of the plant pathogen and biotechnology agent *Agrobacterium tumefaciens* C58. Science. 2001;294(5550):2323–8. Epub 2001/12/18. doi: 10.1126/science.1066803. PMID: 11743194.

36. Curtis PD, Brun YV. Identification of essential alphaproteobacterial genes reveals operational variability in conserved developmental and cell cycle systems. Mol Microbiol. 2014;93(4):713–35. doi: 10.1111/mmi.12686.

37. Vaughan S, Wickstead B, Gull K, Addinall SG. Molecular evolution of FtsZ protein sequences encoded within the genomes of archaea, bacteria, and eukaryota. J Mol Evol. 2004;58(1):19–29. Epub 2004/01/27. doi: 10.1007/s00239-003-2523-5. PMID: 14743312.

38. Margolin W, Long SR. *Rhizobium meliloti* contains a novel second homolog of the cell division gene ftsZ. J Bacteriol. 1994;176(7):2033–43. Epub 1994/04/01. PMID: 8144471.

39. Muller FD, Raschdorf O, Nudelman H, Messerer M, Katzmann E, Plitzko JM, et al. The FtsZ-like protein FtsZm of *Magnetospirillum gryphiswaldense* likely interacts with its generic homolog and is required for biomineralization under nitrate deprivation. J Bacteriol. 2014;196(3):650–9. Epub 2013/11/26. doi: 10.1128/jb.00804-13. PMID: 24272781.

40. Chen Y, Bjornson K, Redick SD, Erickson HP. A rapid fluorescence assay for FtsZ assembly indicates cooperative assembly with a dimer nucleus. Biophys J. 2005;88(1):505–14. Epub 2004/10/12. doi: 10.1529/biophysj.104.044149. PMID: 15475583.

41. Olson BJ, Wang Q, Osteryoung KW. GTP-dependent heteropolymer formation and bundling of chloroplast FtsZ1 and FtsZ2. J Biol Chem. 2010;285(27):20634–43. Epub 2010/04/28. doi: 10.1074/jbc.M110.122614. PMID: 20421292.

42. El-Kafafi el S, Mukherjee S, El-Shami M, Putaux JL, Block MA, Pignot-Paintrand I, et al. The plastid division proteins, FtsZ1 and FtsZ2, differ in their biochemical properties and sub-plastidial localization. Biochem J. 2005;387(Pt 3):669–76. Epub 2004/12/17. doi: 10.1042/bj20041281. PMID: 15601251.

43. TerBush AD, MacCready JS, Chen C, Ducat DC, Osteryoung KW. Conserved dynamics of chloroplast cytoskeletal FtsZ proteins across photosynthetic lineages. Plant Physiol. 2018;176(1):295–306. Epub 2017/08/18. doi: 10.1104/pp.17.00558. PMID: 28814573.

44. Richards DM, Hempel AM, Flardh K, Buttner MJ, Howard M. Mechanistic basis of branch-site selection in filamentous bacteria. PLoS Comput Biol. 2012;8(3):e1002423. Epub 2012/03/17. doi: 10.1371/journal.pcbi.1002423. PMID: 22423220.

45. Howell M, Aliashkevich A, Salisbury AK, Cava F, Bowman GR, Brown PJB. Absence of the polar organizing protein PopZ results in reduced and asymmetric cell division in *Agrobacterium tumefaciens*. J Bacteriol. 2017;199(17). Epub 2017/06/21. doi: 10.1128/jb.00101-17. PMID: 28630123.

46. Ehrle HM, Guidry JT, Iacovetto R, Salisbury AK, Sandidge DJ, Bowman GR. Polar organizing protein PopZ is required for chromosome segregation in *Agrobacterium tumefaciens*. J Bacteriol. 2017;199(17). Epub 2017/06/21. doi: 10.1128/jb.00111-17. PMID: 28630129.

47. Grangeon R, Zupan JR, Anderson-Furgeson J, Zambryski PC. PopZ identifies the new pole, and PodJ identifies the old pole during polar growth in *Agrobacterium tumefaciens*. Proc Natl Acad Sci U S A. 2015;112(37):11666–71. Epub 2015/09/02. doi: 10.1073/pnas.1515544112. PMID: 26324921.

48. Thanbichler M. Synchronization of chromosome dynamics and cell division in bacteria. Cold Spring Harb Perspect Biol. 2010;2(1):a000331. Epub 2010/02/26. doi: 10.1101/cshperspect.a000331. PMID: 20182599.

49. Kuru E, Velocity Hughes H, Brown PJ, Hall E, Tekkam S, Cava F, et al. *In situ* probing of newly synthesized peptidoglycan in live bacteria with fluorescent D-amino acids. Angew Chem Int Ed Engl. 2012;51(50):12519–23. doi: 10.1002/anie.201206749. PMID: 23055266.

50. Alvarez L, Hernandez SB, de Pedro MA, Cava F. Ultra-sensitive, high-resolution liquid chromatography methods for the high-throughput quantitative analysis of bacterial cell wall chemistry and structure. Methods Mol Biol. 2016;1440:11–27. Epub 2016/06/18. doi: 10.1007/978-1-4939-3676-2_2. PMID: 27311661.

51. Szwedziak P, Wang Q, Freund SM, Lowe J. FtsA forms actin-like protofilaments. EMBO J. 2012;31(10):2249–60. Epub 2012/04/05. doi: 10.1038/emboj.2012.76. PMID: 22473211.

52. Ma X, Margolin W. Genetic and functional analyses of the conserved C-terminal core domain of *Escherichia coli* FtsZ. J Bacteriol. 1999;181(24):7531–44. Epub 1999/12/22. PMID: 10601211.

53. Sundararajan K, Goley ED. The intrinsically disordered C-terminal linker of FtsZ regulates protofilament dynamics and superstructure in vitro. J Biol Chem. 2017;292(50):20509–27. Epub 2017/11/02. doi: 10.1074/jbc.M117.809939. PMID: 29089389.

54. Buske PJ, Levin PA. A flexible C-terminal linker is required for proper FtsZ assembly in vitro and cytokinetic ring formation in vivo. Mol Microbiol. 2013;89(2):249–63. Epub 2013/05/23. doi: 10.1111/mmi.12272. PMID: 23692518.

55. Gardner KA, Moore DA, Erickson HP. The C-terminal linker of *Escherichia coli* FtsZ functions as an intrinsically disordered peptide. Mol Microbiol. 2013;89(2):264–75. Epub 2013/05/30. doi: 10.1111/mmi.12279. PMID: 23714328.

56. Khan SR, Gaines J, Roop RM, 2nd, Farrand SK. Broad-host-range expression vectors with tightly regulated promoters and their use to examine the influence of TraR and TraM expression on Ti plasmid quorum sensing. Appl Environ Microbiol. 2008;74(16):5053–62. Epub 2008/07/09. doi: 10.1128/aem.01098-08. PMID: 18606801.

57. Kaczmarczyk A, Vorholt JA, Francez-Charlot A. Cumate-inducible gene expression system for sphingomonads and other alphaproteobacteria. Appl Environ Microbiol. 2013;79(21):6795–802. Epub 2013/09/03. doi: 10.1128/aem.02296-13. PMID: 23995928.

58. Eaton RW. p-Cumate catabolic pathway in *Pseudomonas putida* Fl: cloning and characterization of DNA carrying the cmt operon. J Bacteriol. 1996;178(5):1351–62. Epub 1996/03/01. PMID: 8631713.

59. Wang X, Huang J, Mukherjee A, Cao C, Lutkenhaus J. Analysis of the interaction of FtsZ with itself, GTP, and FtsA. J Bacteriol. 1997;179(17):5551–9. Epub 1997/09/01. PMID: 9287012.

60. Pichoff S, Lutkenhaus J. Identification of a region of FtsA required for interaction with FtsZ. Mol Microbiol. 2007;64(4):1129–38. Epub 2007/05/16. doi: 10.1111/j.1365-2958.2007.05735.x. PMID: 17501933.

61. Moll A, Thanbichler M. FtsN-like proteins are conserved components of the cell division machinery in proteobacteria. Mol Microbiol. 2009;72(4):1037–53. Epub 2009/04/30. doi: 10.1111/j.1365-2958.2009.06706.x. PMID: 19400794.

62. Cameron TA, Zupan JR, Zambryski PC. The essential features and modes of bacterial polar growth. Trends in Microbiology. 2015;23(6):347–53. doi: 10.1016/j.tim.2015.01.003.

63. Meier EL, Razavi S, Inoue T, Goley ED. A novel membrane anchor for FtsZ is linked to cell wall hydrolysis in *Caulobacter crescentus*. Mol Microbiol. 2016;101(2):265–80. Epub 2016/03/31. doi: 10.1111/mmi.13388. PMID: 27028265.

64. Fraipont C, Alexeeva S, Wolf B, van der Ploeg R, Schloesser M, den Blaauwen T, et al. The integral membrane FtsW protein and peptidoglycan synthase PBP3 form a subcomplex in *Escherichia coli*. Microbiology. 2011;157(Pt 1):251–9. Epub 2010/09/18. doi: 10.1099/mic.0.040071-0. PMID: 20847002.

65. Grangeon R, Zupan J, Jeon Y, Zambryski PC. Loss of PopZ At activity in *Agrobacterium tumefaciens* by Deletion or Depletion Leads to Multiple Growth Poles, Minicells, and Growth Defects. MBio. 2017;8(6). Epub 2017/11/16. doi: 10.1128/mBio.01881-17. PMID: 29138309.

66. Rowlett VW, Margolin W. The Min system and other nucleoid-independent regulators of Z ring positioning. Front Microbiol. 2015;6:478. Epub 2015/06/02. doi: 10.3389/fmicb.2015.00478. PMID: 26029202.

67. Monahan LG, Hajduk IV, Blaber SP, Charles IG, Harry EJ. Coordinating bacterial cell division with nutrient availability: a role for glycolysis. MBio. 2014;5(3):e00935–14. Epub 2014/05/16. doi: 10.1128/mBio.00935-14. PMID: 24825009.

68. Flores SA, Howell M, Daniel JJ, Piccolo R, Brown PJB. Absence of the Min system does not cause major cell division defects in *Agrobacterium tumefaciens*. Front Microbiol. 2018;9:681. Epub 2018/04/25. doi: 10.3389/fmicb.2018.00681. PMID: 29686659.

69. Mosyak L, Zhang Y, Glasfeld E, Haney S, Stahl M, Seehra J, et al. The bacterial cell-division protein ZipA and its interaction with an FtsZ fragment revealed by X-ray crystallography. EMBO J. 2000;19(13):3179–91. Epub 2000/07/06. doi: 10.1093/emboj/19.13.3179. PMID: 10880432.

70. Haney SA, Glasfeld E, Hale C, Keeney D, He Z, de Boer P. Genetic analysis of the *Escherichia coli* FtsZ.ZipA interaction in the yeast two-hybrid system. Characterization of FtsZ residues essential for the interactions with ZipA and with FtsA. J Biol Chem. 2001;276(15):11980–7. Epub 2001/03/30. doi: 10.1074/jbc.M009810200. PMID: 11278571.

71. Cho H, Wivagg CN, Kapoor M, Barry Z, Rohs PD, Suh H, et al. Bacterial cell wall biogenesis is mediated by SEDS and PBP polymerase families functioning semi-autonomously. Nat Microbiol. 2016:16172. Epub 2016/09/20. doi: 10.1038/nmicrobiol.2016.172. PMID: 27643381.

72. Meeske AJ, Riley EP, Robins WP, Uehara T, Mekalanos JJ, Kahne D, et al. SEDS proteins are a widespread family of bacterial cell wall polymerases. Nature. 2016;537(7622):634–8. Epub 2016/08/16. doi: 10.1038/nature19331. PMID: 27525505.

73. Emami K, Guyet A, Kawai Y, Devi J, Wu LJ, Allenby N, et al. RodA as the missing glycosyltransferase in *Bacillus subtilis* and antibiotic discovery for the peptidoglycan polymerase pathway. Nat Microbiol. 2017;2:16253. Epub 2017/01/14. doi: 10.1038/nmicrobiol.2016.253. PMID: 28085152.

74. Botta GA, Park JT. Evidence for involvement of penicillin-binding protein 3 in murein synthesis during septation but not during cell elongation. J Bacteriol. 1981;145(1):333–40. Epub 1981/01/01. PMID: 6450748.

75. Morton ER, Fuqua C. Laboratory maintenance of *Agrobacterium*. Curr Protoc Microbiol. 2012;Chapter 1:Unit3D 1. doi: 10.1002/9780471729259.mc03d01s24. PMID: 22307549.

76. Choi YJ, Morel L, Le Francois T, Bourque D, Bourget L, Groleau D, et al. Novel, versatile, and tightly regulated expression system for *Escherichia coli* strains. Appl Environ Microbiol. 2010;76(15):5058–66. Epub 2010/06/22. doi: 10.1128/aem.00413-10. PMID: 20562288.

77. Denkovskiene E, Paskevicius S, Werner S, Gleba Y, Razanskiene A. Inducible expression of *Agrobacterium* Virulence gene VirE2 for stringent regulation of T-DNA transfer in plant transient expression systems. Mol Plant Microbe Interact. 2015;28(11):1247–55. Epub 2015/08/22. doi: 10.1094/mpmi-05-15-0102-r. PMID: 26292850.

78. Morton ER, Fuqua C. Unit 3D.2 genetic manipulation of *Agrobacterium*. Curr Protoc Microbiol. 2012;Chapter:Unit-3D.2. doi: 10.1002/9780471729259.mc03d02s25. PMID: 22549163.

79. Thanbichler M, Iniesta AA, Shapiro L. A comprehensive set of plasmids for vanillate-and xylose-inducible gene expression in *Caulobacter crescentus*. Nucleic Acids Res. 2007;35(20):e137. Epub 2007/10/26. doi: 10.1093/nar/gkm818. PMID: 17959646.

80. Howell M, J. Daniel J, J.B. Brown P. Live cell fluorescence microscopy to observe essential processes during microbial cell growth.:J Vis Exp; 2017.

81. Ducret A, Quardokus EM, Brun YV. MicrobeJ, a tool for high throughput bacterial cell detection and quantitative analysis. Nat Microbiol. 2016;1(7):16077. Epub 2016/08/31. doi: 10.1038/nmicrobiol.2016.77. PMID: 27572972.

82. Lariviere PJ, Szwedziak P, Mahone CR, Lowe J, Goley ED. FzlA, an essential regulator of FtsZ filament curvature, controls constriction rate during *Caulobacter* division. Mol Microbiol. 2018;107(2):180–97. Epub 2017/11/10. doi: 10.1111/mmi.13876. PMID: 29119622.

